# Programmed genome editing of the omega-1 ribonuclease 1 of the blood fluke, *Schistosoma mansoni*

**DOI:** 10.1101/358424

**Authors:** Wannaporn Ittiprasert, Victoria H. Mann, Shannon E. Karinshak, Avril Coghlan, Gabriel Rinaldi, Geetha Sankaranarayanan, Apisit Chaidee, Toshihiko Tanno, Chutima Kumkhaek, Pannathee Prangtaworn, Margaret Mentink-Kane, Christina J. Cochran, Patrick Driguez, Nancy Holroyd, Alan Tracey, Rutchanee Rodpai, Bart Everts, Cornelis H. Hokke, Karl F. Hoffmann, Matthew Berriman, Paul J. Brindley

**Affiliations:** Department of Microbiology, Immunology & Tropical Medicine, & Research Center for Neglected Diseases of Poverty, School of Medicine & Health Sciences, George Washington University, Washington, D.C. 20037, USA; Wellcome Sanger Institute, Wellcome Genome Campus, Hinxton, CB10 1SA, UK; Department of Parasitology, Faculty of Medicine, Khon Kaen University, Khon Kaen, 40002, Thailand; Department of Surgery and the Institute of Human Virology, University of Maryland, Baltimore, MD 21201, USA; Cellular and Molecular Therapeutics Laboratory, National Heart, Lungs and Blood Institute, National Institutes of Health, Bethesda, MD 20814 USA; Department of Parasitology, Faculty of Medicine Siriraj Hospital, Mahidol University, Bangkok, Thailand 10700; Schistosomiasis Resource Center, Biomedical Research Institute, Rockville, MD 20850, USA; Department of Parasitology, Leiden University Medical Center, Leiden, Netherlands; Institute of Biological, Environmental & Rural Sciences (IBERS), Aberystwyth University, Aberystwyth, SY23 3DA, UK

**Keywords:** *Schistosoma mansoni*, genome editing, CRISPR/Cas9, omega1, ribonuclease, flatworms, Platyhelminthes, functional genomics, lentivirus, homology directed repair, non-homologous end joining, double stranded break, Th2 phenotype, monocytic macrophage cell, granuloma, schistosomiasis

## Abstract

CRISPR/Cas9 based genome editing has yet been reported in parasitic or indeed any species of the phylum Platyhelminthes. We tested this approach by targeting omega-1 (ω1) of *Schistosoma mansoni* as a proof of principle. This secreted ribonuclease is crucial for Th2 priming and granuloma formation, providing informative immuno-pathological readouts for programmed genome editing. Schistosome eggs were either exposed to Cas9 complexed with a synthetic guide RNA (sgRNA) complementary to exon 6 of ω1 by electroporation or transduced with pseudotyped lentivirus encoding Cas9 and the sgRNA. Some eggs were also transduced with a single stranded oligodeoxynucleotide donor transgene that encoded six stop codons, flanked by 50 nt-long 5’-and 3’-microhomology arms matching the predicted Cas9-catalyzed double stranded break (DSB) within ω1. CRISPResso analysis of amplicons spanning the DSB revealed ∼4.5% of the reads were mutated by insertions, deletions and/or substitutions, with an efficiency for homology directed repair of 0.19% insertion of the donor transgene. Transcripts encoding ω1 were reduced >80% and lysates of ω1-edited eggs displayed diminished ribonuclease activity indicative that programmed editing mutated the ω1 gene. Whereas lysates of wild type eggs polarized Th2 cytokine responses including IL-4 and IL-5 in human macrophage/T cell co-cultures, diminished levels of the cytokines followed the exposure to lysates of ω1-mutated schistosome eggs. Following injection of schistosome eggs into the tail vein of mice, the volume of pulmonary granulomas surrounding ω1-mutated eggs was 18-fold smaller than wild type eggs. Programmed genome editing was active in schistosomes, Cas9-catalyzed chromosomal breakage was repaired by homology directed repair and/or non-homologous end joining, and mutation of ω1 impeded the capacity of schistosome eggs both to drive Th2 polarization and to provoke formation of pulmonary circumoval granulomas. Knock-out of ω1 and the impaired immunological phenotype showcase the novel application of programmed gene editing in and functional genomics for schistosomes.

## Introduction

Schistosomiasis is considered the most problematic of the human helminth diseases in terms of morbidity and mortality (1-4). The past decade has seen major advances in knowledge and understanding of the pathophysiology, developmental biology, evolutionary relationships and genome annotation of the human schistosomes (5-16). Establishing CRISPR/Cas9 genome editing in schistosomiasis would greatly enable effective functional genomics approaches. The stable CRISPR/Cas9-based site-specific gene mutation and phenotyping will drive innovation and a deeper understanding of schistosome pathogenesis, biology and evolution (17).

The schistosome egg plays a central role both in disease transmission and pathogenesis (1). The appearance of *S. mansoni* eggs in host tissues by six to seven weeks after infection coincides with profound polarization to a granulomatous, T helper type 2 (Th2) cell phenotype (18-22). Numerous egg proteins have been characterized, with >1,000 identified in a well-studied fraction termed soluble egg antigen (SEA) (23-26). In viable eggs, about 30 of the SEA proteins are located outside the developing miracidium and encompass the complement of secreted antigens (egg-secreted proteins, ESP) that interact with host tissues to facilitate the passage of the egg from the mesenteric veins to the intestinal lumen (27). The T2 ribonuclease omega-1 (ω1) is the principal Th2-inducing component of ESP with its Th2-polarizing activity dependent upon both its RNase activity and glycosylation (19, 20, 28). This RNase is hepatotoxic (29), and its secretion by eggs into the granuloma regulates the pattern recognition receptor signaling pathways in dendritic cells that, in turn, prime Th2 responses from CD4^+^ T cells (30). Secreted ω1 provokes granulomatous inflammation around eggs traversing the wall of the intestines, and trapped in hepatic sinusoids and other host organs, driving fibrosis that eventually results in hepatointestinal schistosomiasis (1, 31).

As ω1 drives distinctive immunological phenotypes including Th2 polarization and granuloma formation, we investigated, for the first time in schistosomes and indeed any flatworms, the use of programmed CRISPR/Cas9-mediated genome editing (32, 33) to alter the ω1 locus by both gene knockout and knock-in approaches. The investigation revealed that programmable genome editing catalyzed by the bacterial endonuclease Cas9 was active in schistosomes, with chromosomal double stranded breaks (DSB) repaired by homology directed repair (HDR) using a donor, single stranded oligonucleotide template bearing short homology arms and/or by non-homologous end joining (NHEJ). The programmed mutagenesis decreased levels of ω1 mRNA and induced distinct *in vitro* and *in vivo* phenotypes, including a substantial loss of capacity of SEA from ω1-mutated eggs to polarize Th2 cytokine responses (IL-4 and IL-5) in co-cultured monocytic and T cells and loss of capacity to provoke formation of pulmonary granulomas *in vivo*. Functional knock-out of ω1 and the resulting immunologically impaired phenotype showcase the novel application of CRISPR/Cas9 and its utility for functional genomics in schistosomes.

## Results

### *Omega-1*, a multicopy locus on chromosome 1 of *S. mansoni* specifically expressed in eggs

Five genomic copies of *ω1* were identified in the *S. mansoni* reference genome, version 5 (Fig. s1) although the gene repetition at the ω1 locus on chromosome 1 presents a challenge for genome assembly. A single copy of ω1 selected for genome editing, Smp_193860, included nine exons separated by eight introns and spanned 6,196 nt (Fig. 1a). Several other copies shared similar exon/intron structure and conserved coding sequences (Figs. s1, s2). The predicted coding sequence (CDS) of Smp_193860 encoded a ∼27 kDa protein, of similar mass to the 31 kDa reported for ω1 (34). The gene encodes a secreted ribonuclease of the T2 family of transferase-type endoribonucleases with conserved catalytic regions (35). We designed a sgRNA targeting residues 3808 to 3827 within exon 6 of Smp_193860, adjacent to a AGG protospacer adjacent motif (PAM) and with the predicted Cas9 cleavage site located at three residues upstream of the PAM. (Figure s2 provides the nucleotide sequence of the Smp_193860 copy, and indicates the UTR, coding exons and introns; 6,196 nt.) The AGG and the nucleotide sequence complementary to this sgRNA were also present in the Smp_179960 and Smp_184360 copies of ω1. These three copies shared >99% identity although the CDS of each differed by several substitutions (Fig. s3a-c). All three copies of the *ω1* display a tight profile of developmental stage expression: expression is restricted to the mature egg, and with expression not apparent elsewhere during the developmental cycle of this schistosome (Fig. s1c) (36).

**Figure 1.**
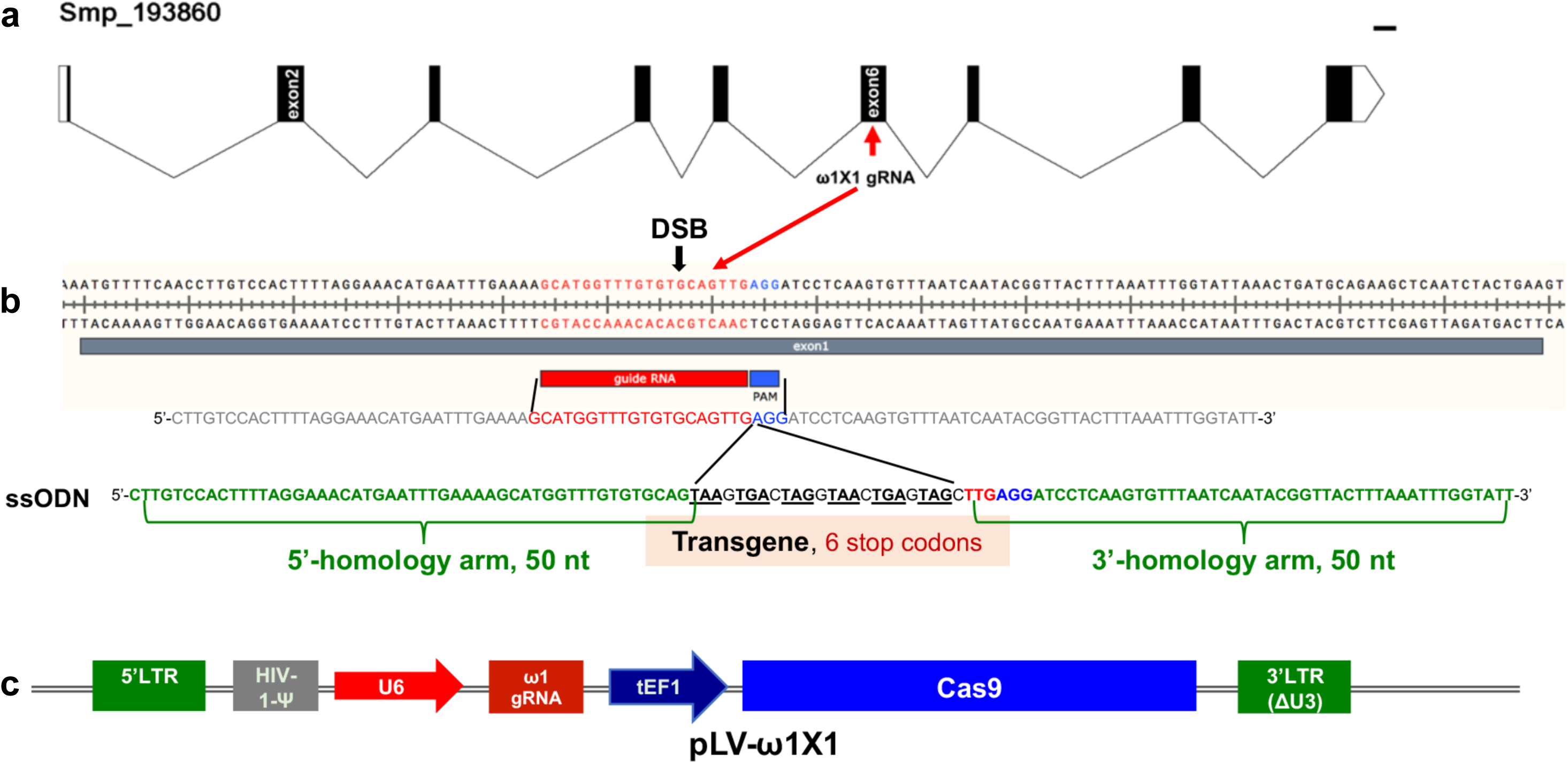
Genomic structure of the locus encoding omega-1 (ω1) in the genome of *Schistosoma mansoni,* guide RNA and CRISPR/Cas9 encoding construct. Gene model of ω1 (*Smp_193860*), showing the position of its nine exons, eight introns and UTRs 6,196 bp on chromosome 1 (**a**). Nucleotide sequence in exon 6 indicating location and sequence of gRNA target site, predicted double stranded break (DSB) (arrow), protospacer adjacent motif (PAM) (AGG, blue box), and 124-nucleotide sequence of the single stranded DNA donor template provided for DSB repair by homologous recombination. Homology arms of 50 nt span a central 24 nt of six-stop-codon transgene (**b**). Linear map of pLV-ω1X1 showing position of regulatory and coding regions for CRISPR/Cas9 editing; the positions of human U6 promoter to drive ω1 gRNA, translational elongation factor EF1-α promoter driving Cas9 from *Streptococcus pyogenes*, and the left and right long terminal repeats of the lentiviral vector derived from HIV-1 (**c**).

### Homology Directed Repair and Non-Homologous End Joining pathways in schistosomes

The draft genome of *Schistosoma mansoni* was surveyed for key proteins of the non-homologous end joining (NHEJ) and homology directed repair (HDR) pathways. Artemis and DNA-PKcs are essential NHEJ factors in vertebrates (37-39). Candidate homologues for six of seven human NHEJ pathway genes and for two key HDR pathway genes, were identified by searching for matches to Pfam (Table s1). A putative homologue of Cernunnos/XLF was not apparent in *S. mansoni* (38) based on searching for the Pfam XLF domain (PF09302) found in human Cernunnos/XLF. The domain appears to be absent from all flatworm species studied by the International Helminth Genomes Consortium (40).

### Site-specific integration of transgene confirmed CRISPR-Cas9 activity in schistosomes

The activity and efficiency of CRISPR/Cas9 to edit the schistosome genome, by targeting the ω1 locus, was explored using two approaches. First, a ribonucleoprotein complex (RNP) comprised of sgRNA mixed with recombinant Cas9 endonuclease was delivered into schistosome eggs isolated from livers of experimentally-infected mice (eggs termed ‘LE’, ‘liver eggs’) by electroporation. In addition, Homology Directed Repair (HDR) of CRISPR/Cas9-induced double stranded breaks (DSBs) at the ω1 locus in the presence of a donor DNA template was investigated (41-43). A single stranded oligodeoxynucleotide (ssODN) of 124 nt in length was delivered to some LE as a template for HDR of chromosomal DSBs (Fig. 1b). The ssODN included a short transgene encoding six stop codons flanked by 5’-and 3’-homology arms, each arm 50 nt in length, complementary to the genome sequence of exon 6 on the 5´ and 3´ sides of the Cas9 cleavage site (Figs. 1a, 1b). In a second approach, a lentivirus vector (pLV-ω1X1; Fig. 1c) that included Cas9, driven by the mammalian translational elongation factor 1 promoter, and the exon 6-targetting sgRNA (20 nt), driven by the human U6 promoter was engineered (44). LE were transduced with pseudotyped lentiviral virions (LV) by exposure in culture to LV for 24 hours (45) and, thereafter, transfected with the ssODN repair template. In both approaches, expression of ω1 in LE after 72 hours in culture was ascertained.

Given that the donor ssODN included a short transgene that facilitates genotyping, PCR was performed using template genomic DNAs from the CRISPR/Cas9-treated LE (41) to reveal the site-specific knock-in (KI). A forward primer termed Smω1X1-6stp-cds-F specific for the ssODN transgene was paired with three discrete reverse primers, termed Smω1-R1, Smω1-R2 and Smω1-R3, at increasing distance from the predicted HDR insertion site in ω1 (Table s2). Amplicons of the expected sizes of 184, 285 and 321 nt were observed in genome-edited eggs but not in control eggs (Fig. 2a, b; Fig. s4a), a diagnostic pattern indicative of the ssODN transgene insertion into ω1 and, in turn, indicating that the resolution of the DSB at the ω1 locus from CRISPR/Cas9 had been mediated by HDR. Amplification using a control primer pair that spanned the predicted DSB, termed Smω1-control-F/R, yielded control amplicons of the expected 991 nt. Similar findings were observed with genome editing delivered by RNPs and by LV (Fig. s4). Sanger sequence analysis of the knocked-in amplicons (KI-R1, KI-R2 and KI-R3) confirmed the presence of the transgene inserted into the ω1 locus at the site predicted for programmed Cas9 cleavage (Fig. 2c).

**Figure 2.**
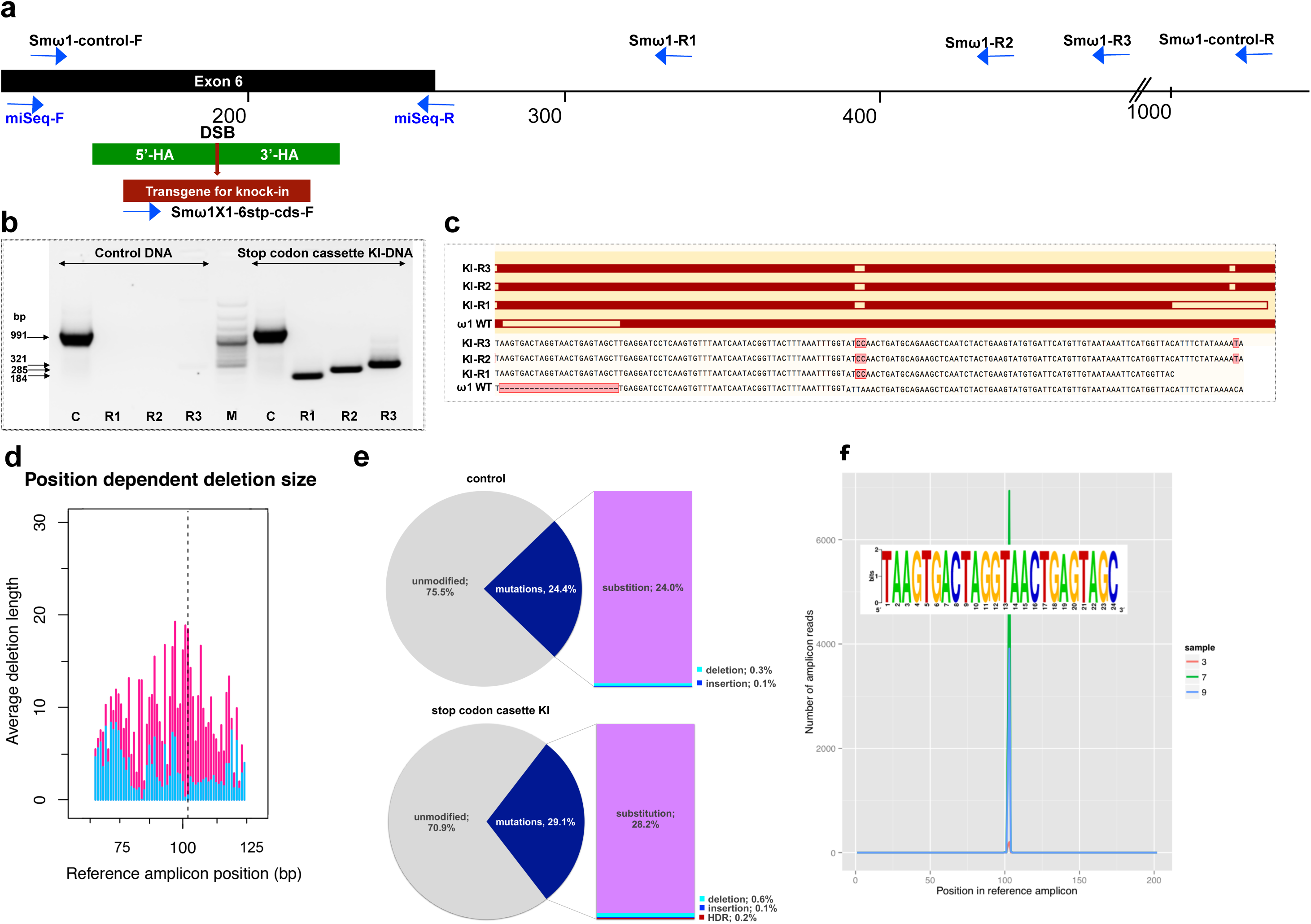
Programmed chromosomal break at ω1 locus repaired by homologous recombination from donor template. Schematic diagram to indicate positions of primer binding sites (blue arrows), with the foreign gene cassette as the forward primer (Smω1X1-6-stop codons cassette-F) paired with three discrete reverse primers, Smω1-R1, Smω1-R2 and Smω1-R3 from the ω1 locus and a primer pair for target amplicon NGS library amplification; miSeq-F and miSeq-R (**a**). The control PCR amplicon was generated using the Smωω1-control-F and –R primers. The green box shows the location of 5’ and 3’ homology arms, the red box and arrow indicate the stop codon bearing transgene. **b**, PCR products visualized in ethidium bromide-stained agarose gel demonstrating Cas9-catalyzed target site-specific insertional mutagenesis in exon 6 of the ω1 gene. Evidence for transgene knocked-in into programmed target site revealed by amplicons of the expected sizes in lanes R1, R2 and R3, of 184, 285 and 321 bp, respectively (arrows at left) spanned the mutated site in the genomic DNAs pooled from schistosome eggs, including a positive control flanking the insert site (991 bp). The control DNA result shown in this gel was isolated from heat-inactivated-pLV-ω1X1 virions and ssODN treated LE. Similar findings were obtained when programmed gene editing was executed by lentiviral virion-delivered Cas9 and ω1-gRNA transgenes and by ribonucleoprotein complex (RNP) delivered by square wave electroporation (supporting information). The non-KI control groups (sgRNA only, heat-inactivated pLV-ω1X1 virions only, ssODN only) showed no amplicons by stop cassette-KI primers with R1, R2 or R3 (Fig. s3). **c**, Multiple sequence alignments confirmed the presence of the 24 nt transgene inserted precisely into exon 6 of ω1 locus from KI-R1, -R2 and -R3 fragments compared with ω1 wild type (WT). The white box on ω1-WT indicates the absence of the transgene sequence and white boxes on KI-R1, -R2 and –R3 fragments show locations of substitutions relative to the other ω1 copies (Smp_184360): 2 bp (AT to CC) mismatches at positions 253-254 nt. All three contained the (knock-in) insertion sequence (white box), which confirmed targeted mutation of the ω1 gene. **d-f**, Illumina deep sequence analysis of amplicon libraries revealed Cas9 induced on-target repair of programmed gene mutation of the ω1 locus by deletions, insertions, and substitutions by CRISPResso analysis. **d**, Frequency distribution of position-dependent deletions and of deletion sizes; these varied from point mutations to >20 bp adjacent to the DSB. The dotted line indicates the predicted position of the programmed double stranded break. **e.** Frequency of frameshift versus in-frame mutations reported by CRISPResso. The pie charts show the fraction of all mutations (indels and substitutions) in the coding region (positions 42-179) of the amplicon predicted to induce frameshifts, *i.e.* indels of 1-2 bp, or multiples thereof. Top graph corresponds to sample 2 (eggs only control) (Table s3), bottom graph corresponds to sample 9 (eggs exposed to virions and ssODN, i.e. CRISPR/Cas9-treated) (Table s3). Findings for control and treated samples are provided in Table s3. **f.** Insertions of the knock-in sequence. Number of amplicon reads containing an insertion of the knock-in sequence (with ≥75% identity to it) is shown in the Y-axis, and the position of the insertion relative to the reference amplicon is shown on the X-axis. The programmed Cas9 scission lies between positions 102 and 103. Samples 3, 7 and 9 are independent amplicon libraries (technical replicates) made from the same sample of genomic DNAS pooled from six biological replicates exposed to virions and ssODN. The insert shows a sequence logo, created using WebLogo (94), of the sequences of the 3,826 sequence reads from samples 3, 7 and 9, with insertions of 24 bp at position 102; most matched the donor template, <u>TAA</u>G<u>TGA</u>C<u>TAG</u>G<u>TAA</u>C<u>TGA</u>G<u>TAG</u>C.

### Programmed mutations in exon 6 of ω1

The activity of CRISPR/Cas9 was first evaluated by a quantitative PCR (qPCR) approach that relies on the inefficient binding of a primer overlapping the gRNA target, i.e. where mutations are expected to have occurred, compared to the binding efficiency of flanking primers, i.e. outside the mutated region (46, 47). The overlapping (OVR) primer pair shared the reverse primer with OUT primer (Fig. s5a). Genomic DNA template was used for qPCR to quantify the efficiency of CRISPR-mediated mutagenesis at the target locus; the ratio between the OVR products and OUT products estimate the relative fold amplification reduction in CRISPR/Cas9-manipulated samples compared to controls in the target sequence of the gRNA, i.e. the annealing site for the OVR primer. Relative fold amplification was reduced by 12.5% in gDNA isolated from eggs treated with pLV-ω1X1 and ssODN whereas a reduction in relative fold amplification of 2.5, 6.9, and 4.5 were observed in eggs treated with gRNA/Cas9 RNP complex alone, gRNA/Cas9 RNP complex and ssODN, or pLV-ω1X1 alone, respectively. A reduction in relative fold amplification was not apparent among control groups, i.e. untreated eggs, eggs electroporated in the presence of Opti-MEM only, Cas 9 only, eggs transduced with heat-inactivated pLV-ω1X1 with ssODN donor, and eggs transfected with ssODN only (Fig. s5b).

To further characterize and quantify the mutations that arose in the genome of *ω1* gene-edited eggs, we used an amplicon-sequencing approach. Barcoded amplicon libraries were constructed from pooled genomic DNA of six independent exposures of LE to pLV-ω1X1 and the donor ssODN. Each amplicon was sequenced on the MiSeq Illumina platform and the CRISPResso pipeline (48) was used to analyze deep-coverage sequence. More than 56 million sequenced reads were compared to the reference sequence of the Smp_193860 locus (Table s3), which revealed that 71% exhibited the wild type (WT; i.e., unmodified DNA) whereas 29 % reads exhibited apparent mutations (Fig. 2d) across the 202 bp amplicon, with 0.13% insertions, 0.58% deletions and 28.2% substitutions (Table s3, sample 9). In contrast, in the control eggs-only group, 76% were WT, and 24% of reads exhibited apparent mutations, with 0.14% insertions, 0.33% deletions, and 24.0% substitutions (Fig. 2e, sample 2 in Table s3). Thus, subtracting the rate of apparent mutations in the control, we estimated that 0.25% and 4.2% of reads in the experimental sample carried programmed CRISPR-induced deletions and substitutions, respectively. Indels of 1-2 bp, or multiples thereof, in coding DNA cause frame-shifts, and consistent with its higher rate of indels, the CRISPR/Cas9-treated sample displayed a higher rate of frame-shifts compared to a sample from control eggs, 2.0 % versus 1.4 %) (Table s3).

Many apparent sequence variants common to the control and edited eggs likely reflect polymorphism among copies of ω1 rather than programmed mutations. The sequence reads revealed several common variants, such as adjacent ‘TA’ substitutions instead of ‘CC’ at positions 152-153 of the amplicon, which encodes a change from Q to K at the amino acid level. The gene Smp_193860 has ‘TA’ at this position in the V5 assembly (12), as does the mRNA XP_018647487.1 from the NCBI database, whereas Smp_193860, Smp_184360 and Smp_179960 all have ‘CC’ at this position in the V7 assembly (Berriman and co-workers, in preparation) (Fig. s3 a, b), whereas ‘CC’ was also observed in KI fragments by Sanger direct sequencing (Fig. 2c). A second common dinucleotide substitution from ‘AC’ to ‘TT’ at positions 60-61 encodes an amino acid change from T to F. Both dinucleotide substitutions occurred together in 8% of reads in the control group (Table s3, sample 2,) and 4% of reads in the gene-edited group (Table s3, sample 9). These non-synonymous substitutions may have functional significance given their proximity to the catalytic site of the ribonuclease (Fig. s3c).

Along with the predicted NHEJ-catalyzed mutations, CRISPResso determined the rate of HDR-mediated ssODN knock-in (Table s3; Fig. 2f). Here, insertion of the 24 bp transgene was confirmed in 0.19% of the reads at the sgRNA programmed CRISPR/Cas9 target site (Fig. 2f; sample 9 in Table s3). Some reads containing the knock-in sequence included the ‘CC’ to ‘TA’ substitutions at positions 152-153 and ‘AC’ to ‘TT’ at positions 60-61 (Fig. s3; Table s3). This indicates that the indels catalyzed by NHEJ and/or KI by HDR occurred in multiple copies of ω1 including Smp_193860, Smp_184360 and Smp_179960, and possibly also further copies not yet annotated in the reference genome. The qPCR approach estimated a reduction by 12.5% in relative fold amplification in the pLV-ω1X1 with ssODN treatment group (Fig. s5) whereas CRISPResso analysis of the pooled NGS reads indicated a frequency of indel/substitution mutation of ∼4.5% (Table s3).

### Programmed gene editing markedly diminished the expression of ω1

Liver eggs (LE) transfected with RNP complexes, without or with ssODN, displayed a down regulation of the ω1-specific transcript of ∼45% and 81%, respectively, compared to controls (*p* ≤ 0.05; *n* = 11). However, LE transduced with pLV-ω1X1 virions, without or with ssODN, showed a reduction of the *ω1*-specific transcripts of 67% and 83% respectively, when compared to controls (Fig. 3a). Similar outcomes were seen in all biological replicates undertaken (*n* =11). This outcome indicated that resolution of chromosomal DSB by NHEJ plus HDR provided enhanced programmed gene knockout compared to NHEJ-mediated chromosomal repair alone. Nevertheless, both RNPs and pLV virions efficiently delivered programmed gene editing to schistosomes but lentiviral transduction delivered enhanced performance with stronger gene silencing, in particular when the donor repair template was provided (Fig. 3a). When examined at later time points (days 5 and 7 following manipulation of the LE), further reduction in ω1 abundance was not apparent (Fig. 3b).

**Figure 3.**
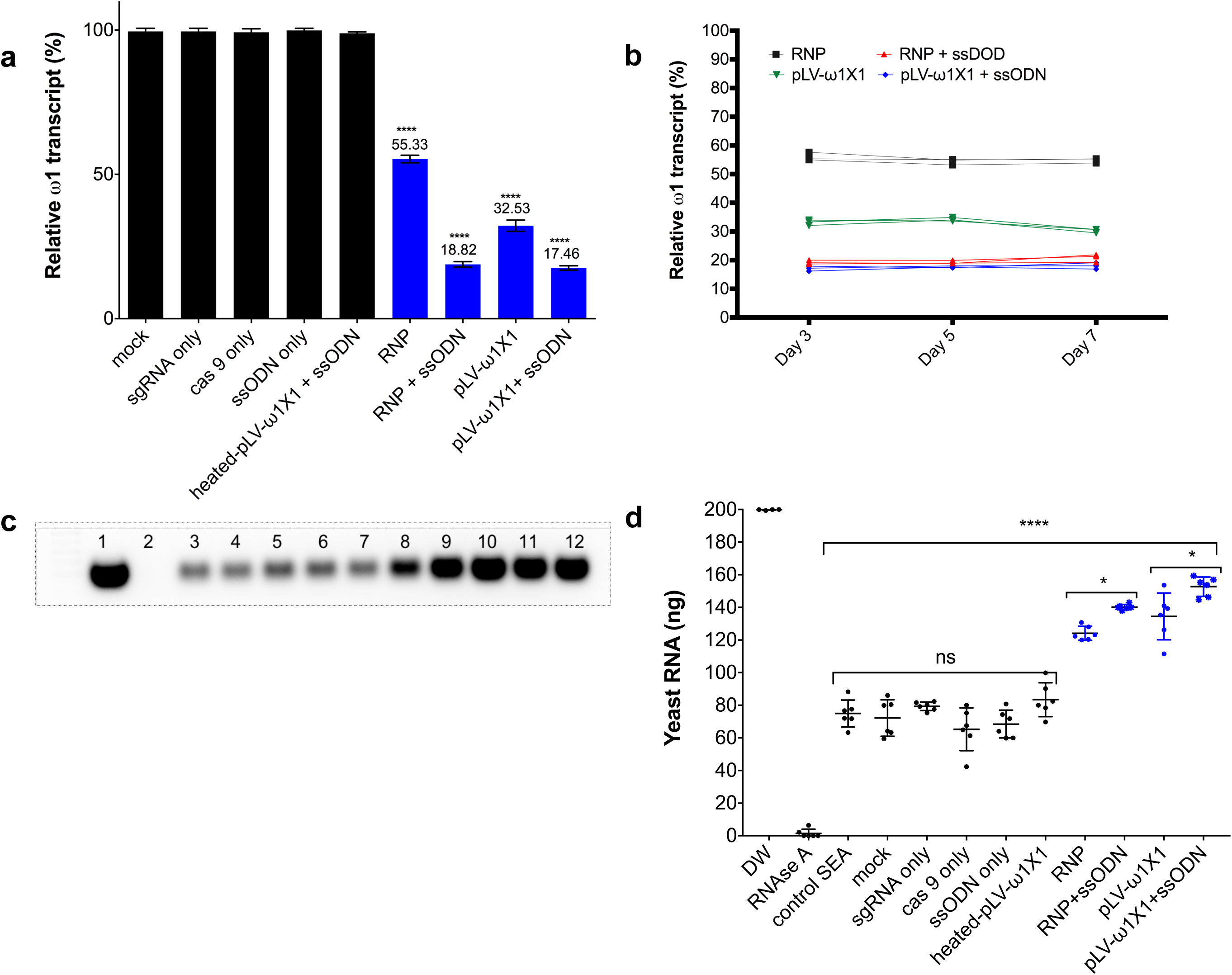
Diminished ω1-specific transcript levels and ribonuclease T2 activity following programmed editing. Panel **a,** ω1 mRNA abundance was reduced up to ∼ 70% after genome editing by sgRNA/Cas9 complex and lentivirus systems, and markedly reduced >80% with the addition of ssODN as the DNA repair donor. Relative expression of ω*1* transcripts at three days following CRISPR/Cas9 manipulation; mean ±SD, *n* =11 (biological replicates); *p* ≤ 0.0001 (****) (ANOVA). **b,** Stable reduction of ω*1* transcripts at days 5 and 7 after treatment (three biological replicates) in four experimental groups; RNP (black), RNP and ssODN (red), pLV-

Large DNA deletions have been associated with CRISPR/Cas9 mutations in another helminth species (42). However, using qPCR to estimate relative copy number, as previously described (49), evidence that silencing of *ω1* was associated with a reduction in the copy number of this multi-gene locus was not seen (Fig. s6).

### Diminished ribonuclease activity in CRISPR/Cas9-mutated schistosome eggs

The ribonuclease activity of the ω1 glycoprotein in SEA is associated with the Th2-polarized immune response that underpins the appearance of schistosome egg granulomata (19, 22). Ribonuclease activity of SEA from control and experimental groups on substrate yeast RNA was investigated following CRISPR/Cas9 programmed mutation of ω1 mediated by the RNP and the pseudotyped lentiviral approaches with or without ssODN (20, 50). Intact yeast RNA was evident in the DNase-RNase free condition (negative control), indicative of absence of RNase activity in the reagents (200 ng yeast RNA at the outset). There was near complete hydrolysis of yeast RNA following exposure to RNase A (positive control); ∼1.4 ng of RNA remained intact. Wild type SEA exhibited marked RNase activity against the yeast RNA; ∼70 ng RNA remained intact after one hour, corresponding to >60% digestion. Incubation of the RNA with Δω1-SEA from the experimental groups, RNP, RNP + ssODN, pLV-ω1X1, and pLV-ω1X1+ssODN, resulted in ∼30% substrate digestion, with 124, 140, 135 and 153 ng of RNA remaining, respectively. All conditions for programmed genome editing resulted in less digestion of the yeast RNA than for wild type SEA (*p* ≤ 0.0001) (Figs. 3c, 3d). Moreover, the Δω1-SEA with programmed knock-in exhibited less RNase activity than Δω1-SEA prepared without the donor ssODN repair template (*p* ≤ 0.01).

### Depleting SEA of ω1 down-regulated Th2 response

The ω1 ribonuclease alone is capable of conditioning human monocyte-derived dendritic cells to drive Th2 polarization (22) and enhanced CD11b^+^ macrophage modulation of intracellular toll like receptor (TLR) signaling (30). Ribonuclease ω1 inhibited TLR-induced production of IL-1β and redirected the TLR signaling outcome towards an anti-inflammatory posture via the mannose receptor (MR) and dectin (22, 51, 52). The human monocytic cell line, THP-1, and the Jurkat human CD4^+^ T cell line were employed to investigate the interaction of antigen presenting cells and T cells (53, 54). At the outset, the THP-1 cells were differentiated to macrophages for 48 hours, then pulsed with SEA or Δω1–SEA for 48 hours, after which the Jurkat CD4^+^ T cells were added to the wells. Subsequently, the co-culture continued for 72 hours. Representative cytokines, including IL-4, IL-5, IL-13, IL-2, IL-6, IL-10, TNF-α and IFN-γ, were quantified in supernatants of the co-cultures (Fig. 4). SEA from ω1-mutated eggs induced reduced levels of Th2 cytokines, including IL-4 and IL-5, in comparison to wild type SEA (*p* ≤ 0.01), and a trend towards less IL-13 production was also observed (Fig. 4). Moreover, reduced levels of IL-6 and TNF-α were observed (*p* ≤ 0.01). By contrast, significant differences in levels of IL-10 and IL-2 were not evident between the WT-and mutant-SEA groups. IFN-γ was not detected following pulsing with the WT-SEA or mutant-SEA (Fig. s7).

**Figure 4.**
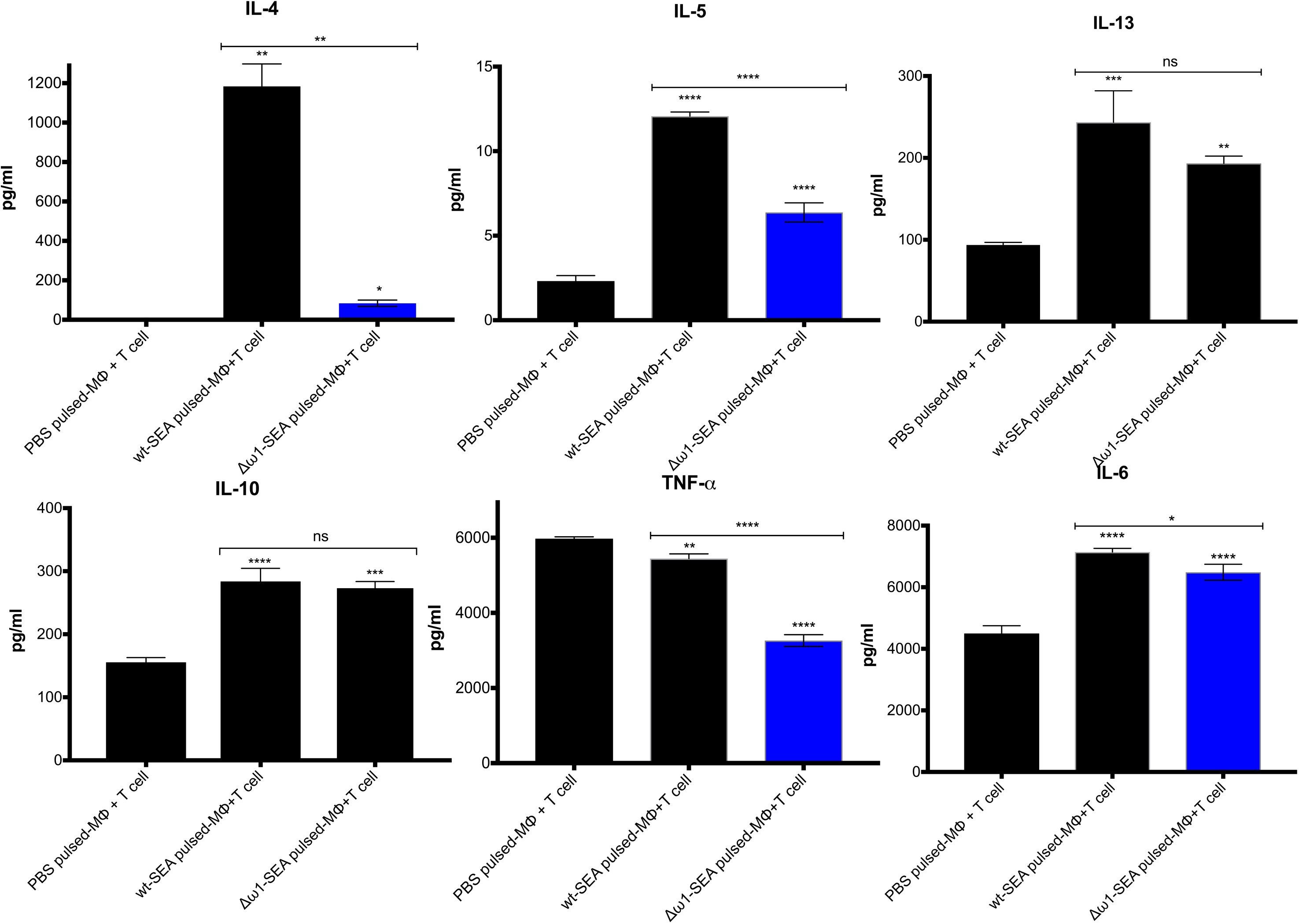
Reduced Th2 cytokine levels following exposure to Δω1-SEA. Reduction in Th2 cytokines IL-4 and IL-5 but not IL-13 (95, 96) following pulsing of Mϕ (PMA induced-THP-1 cells) with Δω1-SEA prior to co-culture with human CD4^+^ T cells (Jurkat cell line) compared with WT-SEA pulsed-Mϕ (top panels). In addition, levels of IL-6 and TNF-α were reduced where Mϕ were first pulsed with Δω1-SEA but not WT SEA. Differences were not evident for IL-10. The assay was carried out in triplicate; *p* < 0.0001, ≤ 0.0001, 0.0038 and 0.0252 indicated as ****, ***, ** and *, respectively (one-way ANOVA, multiple comparison).

### Granulomatous inflammation markedly reduced in lungs of mice injected with Δω1 eggs

Following the entrapment of eggs in the intestines, liver and eventually lungs, the glycosylated ω1 ribonuclease represents the principal stimulus that provokes the development of the circumoval granuloma, necessary for extravasation of the eggs (55). A long-established model of the schistosome egg granuloma employs tail vein injection of eggs into mice, which leads to formation of circumoval granuloma in the mouse lung (56-58). The latter approach has been extensively employed for immunopathogenesis-related studies of ω1 (16). Accordingly, to elicit circumoval granulomas, ∼3,000 WT or Δω1 LE were injected into the lateral vein of the tail of BALB/c mice. The mice were euthanized 10 days after injection, and the entire left lung was removed, fixed, sectioned, and stained for histological analysis (Fig. 5). Representative digital microscopic images of the whole mouse lungs acquired through high-resolution 2D digital scans are presented in Figure 5, panels a-g. At low magnification (2×), much more severe and widespread inflammation was visible in lungs exposed to WT eggs compared to Δω1-eggs. In addition, markedly more intense and dense focal inflammation was induced by WT compared to Δω1-eggs (Fig. 5b). Granulomas were not seen in control naïve mice not exposed to schistosome eggs (Fig. 5c). At 20× magnification, vast disparity in volume of the circumoval granulomas was observed for WT versus Δω1 LE (Fig. 5a1-a2, 5d, 5e *vs.* Figs. 5b1-b2, 5f, 5g). The volume of granulomas surrounding single schistosome eggs was quantified; those surrounding WT eggs in lungs of the mice were 18-fold greater than for Δω1-eggs, 21×10^−2^±1.61×10^−3^ mm^3^ and 0.34×10^−2^ ±0.12×10^−4^ mm^3^ (mean ± S.E., 17-26 granulomas per mouse), respectively (*p* < 0.0001) (Fig. 5h). The experiment was repeated with 3-4 mice per group with a similar outcome. The findings documented marked deficiency in the induction of pulmonary granulomas by the Δω1 compared to WT eggs of *S. mansoni*.

**Figure 5.**
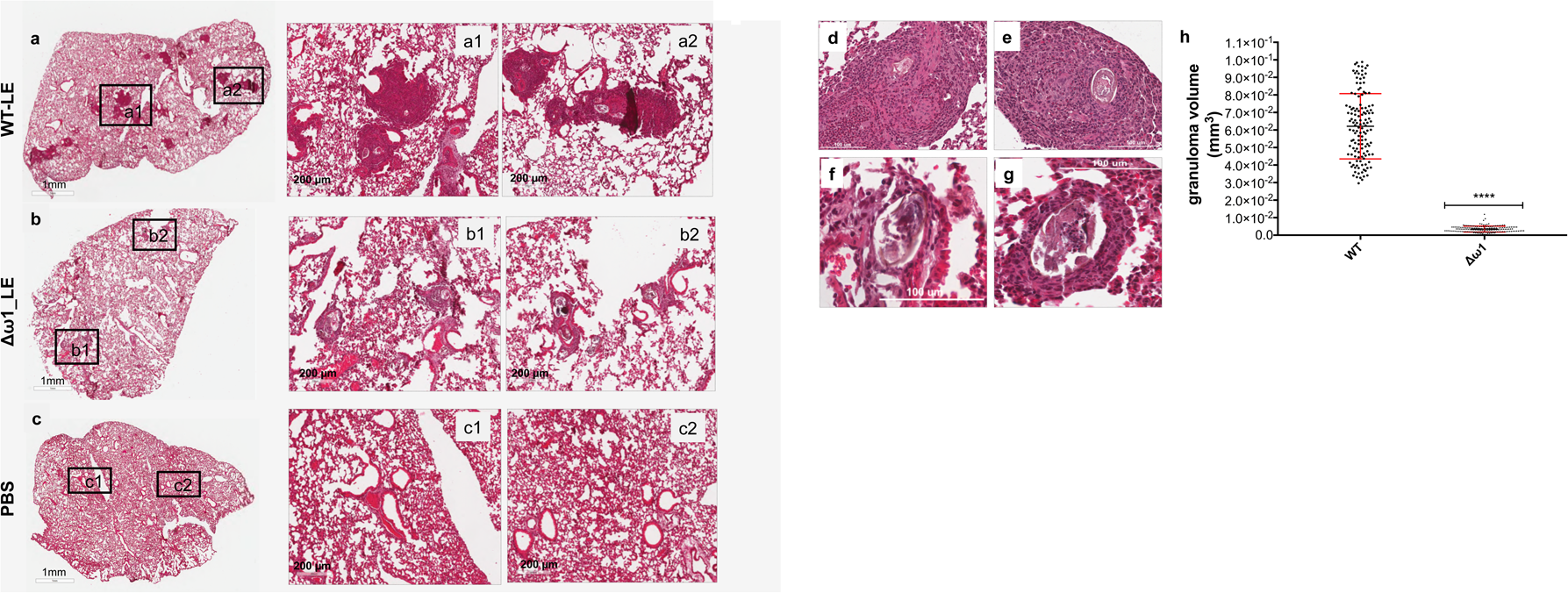
Pulmonary circumoval granulomas revealed attenuated granulomatous response to Δ. ω1 **schistosome eggs.** Schistosome eggs (∼3,000 eggs) that had been transduced with lentivirus virions encoding ω1-specific sgRNA and Cas9 in tandem with ssODN were introduced via the tail vein into mice. The mice were euthanized 10 days later; thin sections of the left lung were stained with H&E, and circumoval granulomas counted and measured. Representative 2D scanned micrographs of granulomas inoculated with WT eggs (**a**) (2× magnification) and 20× magnification (a1 and a2), and with Δω1 eggs; **b**, (2×), b1 and b2 (20×). **c**, Control mouse lung. **d, e**, Representative micrograph of individual, control egg induced-granuloma that were counted to assess for granuloma volume. **f, g**, Representative micrographs showing Δω1 egg induced-granulomas. All single egg induced-granuloma from WT and Δω1 eggs were measured and granuloma volume estimated (25). **h,** Scatter plots of the volume (mm^3^) for individual granuloma, mean ± SE (red) is shown. Granulomas induced by Δω1 eggs were significantly smaller than those surrounding WT eggs (Welch’s *t*-test, *p* ≤ 0.0001, *n* >100).

### GenBank/EMBL/DDBJ

***Database accessions*** Sequence reads from the amplicon NGC libraries are available at the European Nucleotide Archive, study accession number ERP110149. Additional information is available at Bioproject PRJNA415471, https://www.ncbi.nlm.nih.gov/bioproject/PRJNA415471 and GenBank accessions SRR6374209, SRR6374210.

## Discussion

Using *Schistosoma mansoni* as an exemplar, we have demonstrated the activity and feasibility of programmed CRISPR/Cas9 genome editing in the phylum Platyhelminthes. On-target genome editing was evidenced by site-specific mutations at the *ω1* locus on chromosome 1. The chromosomal lesion was repaired by the non-homologous end joining (NHEJ) pathway (59) in the absence of a donor oligonucleotide and by homology directed repair (HDR) when a single stranded oligonucleotide donor template was provided (60-62).

To investigate the feasibility of genome editing, schistosome eggs were transfected with ribonucleoprotein particle (RNP) complexes and with lentiviral virions carrying the CRISPR/Cas9 components, in similar fashion to earlier reports in cell lines, tissues and entire organisms (63-66). Delivery by RNPs facilitates immediate editing although the short half-life of the RNP components may be disadvantageous. Delivery by plasmids and by viral-mediated infection may provide sustained Cas9 activity, transgene integration in non-dividing cells, and other advantages (33, 67). Transfection by LV provides a hands-free approach to enable scaling of gene editing and the potential that less accessible and/or differentiated cells can be reached. When released into the mesenteric veins from adult female, the schistosome ovum each contains single celled zygote surrounded by 30 to 40 vitelline cells (27). By six days later, the miracidium that has developed within the eggshell is a multi-cellular, mobile, ciliated larva comprised of organs, tissues, muscles and nerves (25, 68-70).

To investigate the efficiency of programmed genome editing, several parallel approaches were undertaken including NGS-and quantitative PCR-based analysis of pooled genomic DNAs, analysis of levels of ω1-transcripts and ribonuclease, and immuno-phenotypic status of cultured cells and of mice exposed to gene edited eggs. Analysis of the deep-coverage nucleotide sequence reads of amplicons spanning the predicted DSB site in the *ω1* locus revealed that ∼4.5% of the reads were mutated by insertions, deletions, and substitutions. The target locus was mutated by knock-in (KI) of a ssODN repair template bearing short homology arms to mediate homology directed repair following DSB at *ω1*, with an efficiency for HDR of 0.19% insertion of the donor transgene. Numerous substitutions in addition to deletions and insertions were seen, although some may reflect single nucleotide polymorphisms (SNPs) among the multiple gene copies rather than genome editing-induced changes. As well as the anticipated ease of access by the RNPs, virions and the donor template ssODN to cells proximal to the eggshell compared to the cells deeper within the egg, other factors may have contributed to variability of CRISPR efficiency among the eggs and the cells within individual eggs. Given the multiple copies of *ω1*, the numerous cells in the mature egg (68), the spectrum of development of the eggs in LE, and other factors, a mosaic of mutations would be expected: some genomes to display HDR but not NHEJ, some NHEJ but not HDR, others both NHEJ and HDR, and many to retain the wild type genotype. HDR proceeds at cell division, with the cell at late S or G2 phase following DNA replication where the sister chromatid serves as the repair template; otherwise, NHEJ proceeds to repair the DSB (71).

Expression levels of *ω1* were diminished by as much as 83% relative to controls suggesting that Cas9 catalyzed the mutation of *ω1*, and that programmed Cas9-induced DSBs had been resolved by NHEJ and/or HDR. Knock-in of the ssODN repair template induced 81-83 % reduction in ω1-specific mRNA levels, whereas down regulation of 45 to 67 % followed the exposure of eggs to RNP or lentivirus without ssODN. Ribonuclease activity in *ω1*-mutated eggs was likewise significantly diminished. Curiously, less than 5% efficiency (NGS findings) in gene editing appeared to account for this markedly reduced (>80%) gene expression. The possibility of large-scale deletions, as reported in *Strongyloides stercoralis* (42), presented an explanation for this apparent paradox. However, analysis of copy number by qPCR failed to reveal apparent differences for *ω1* among treatment and control groups of eggs. An alternative explanation for the paradox may be the tight stage-and tissue-specific expression of ω1. Within the mature egg, the fully developed miracidium is surrounded by a squamous, syncytial epithelium termed variously as the envelope, the inner envelope, or von Lichtenberg’s envelope (25, 27, 68, 69). This layer is metabolically active (27) and is considered to be the site of synthesis of the ω1 T2 ribonuclease that is released from the egg into the granuloma (25, 27, 29), along with other secreted/ excreted proteins that facilitate egress of the egg from the blood vessels and through the wall of the intestine (27). Expression of ω1 is developmentally tightly regulated: based on comparison of expression in the egg to expression in the miracidium, immunostaining and transmission electron microscopy of the mature egg, and on meta-analysis of transcriptomic findings, all or most expression of ω1 occurs solely in the mature egg; expression does not occur in the immature egg or other developmental stages (20, 25, 27, 29, 72, 73) (Fig. s1c). The virions may have transduced only a small number of the cells within the mature egg because, after entry into the egg, the virus would be expected to contact the inner envelope, rather than traveling further to the cells of the miracidium. Accordingly, the efficiency of gene editing may have ranged from high to low through the cells of the egg due to ease of virus access, and displayed highest efficiency in the cells of the inner envelope where ω1 is expressed. This may explain the paradox involving >80% reduction in ω1-encoding transcripts in tandem with the <5% gene editing efficiency.

To address the effects on the hallmark Th2 polarization of the immune response to schistosomiasis (74), co-cultures of human macrophages and T cells were exposed to Δω1-SEA and mice were exposed to Δω1-eggs. Whereas wild type SEA polarized Th2 cytokine responses including IL-4 and IL-5 in the co-cultures, significantly reduced levels of these cytokines were observed after exposure to ω1-mutated SEA. Moreover, following introduction of eggs into the tail vein of mice, the volume of pulmonary circumoval granulomas around Δω1 eggs was enormously reduced compared to those provoked by wild type eggs. Whereas this outcome extends earlier findings using lentiviral transduction of eggs of *S. mansoni* to deliver microRNA-adapted short hairpin RNAs aiming to silence expression of ω1 (16), the decrement in granuloma volume was vastly more marked in the present study. In addition to incomplete disruption of all copies of ω1, residual granulomas around the mutant eggs may be due to the presence of other Th2-polarizing components within SEA that have recently been reported (75). Given that the T2 ribonuclease ω1 is the major type 2-polarizing protein among egg-secreted proteins (19, 20), our findings of a phenotype characterized by the absence of or diminutive granulomas provide functional genomics support to this earlier advance (22).

## Conclusions

These findings confirmed that somatic genome editing of schistosome eggs led to functional knockout of the ω1 T2 ribonuclease, and mosaicism of mutant and wild type cells. The genome-edited eggs exhibited loss of function of ω1 but remained viable. Programmed mutation of ω1 using CRISPR/Cas9 not only achieved the aim of establishing the applicability of genome editing for functional genomics of schistosomes but also demonstrated manipulation of a gene expressed in the schistosome egg, the developmental stage central to the pathophysiology of schistosomiasis. This study provides a blueprint for editing other schistosome genes, and those of trematodes and platyhelminths at large. The challenge for future studies is now to deliver pseudotyped virions and programmed genome editing to the schistosome germ line. Mutant parasite lines derived following this protocol (11, 76) will enable more comprehensive understanding of the pathogenesis of this neglected tropical disease and accelerate the discovery of novel strategies for parasite control.

## Materials and Methods

### Ethics statement

Mice experimentally infected with *S. mansoni*, obtained from the Biomedical Research Institute, Rockville (BRI), MD were housed at the Animal Research Facility of the George Washington University Medical School, which is accredited by the American Association for Accreditation of Laboratory Animal Care (AAALAC no. 000347) and has an Animal Welfare Assurance on file with the National Institutes of Health, Office of Laboratory Animal Welfare, OLAW assurance number A3205-01. All procedures employed were consistent with the Guide for the Care and Use of Laboratory Animals. The Institutional Animal Care and Use Committee of the George Washington University approved the protocol used for maintenance of mice and recovery of schistosomes. Studies with BALB/c mice involving tail vein injection of schistosome eggs and subsequent euthanasia using overdose of sodium pentobarbitone was approved by the IACUC of the Biomedical Research Institute (BRI), protocol 18-04, AAALAC no. 000779 and OLAW no. A3080-01.

### Schistosome eggs

Mice were euthanized seven weeks after infection with *S. mansoni*, livers were removed at necropsy, and schistosome eggs recovered from the livers, as described (77). The liver eggs termed ‘LE’ were maintained in DMEM medium supplemented with 10% heat-inactivated fetal bovine serum (FBS), 2% streptomycin/penicillin at 37°C under 5% CO_2_ in air (76, 78). Polymyxin B to (10 μg/ml) was added to the cultures twice daily to neutralize lipopolysaccharide (LPS) (79). Soluble egg antigen (SEA) was prepared from these eggs, as described (23, 56). In brief, the homogenate of eggs in 1×PBS containing protease inhibitor cocktail (Sigma) was frozen and thawed twice, clarified by centrifugation at 13,000 rpm, 15 min, 4°C, the supernatant passed through a 0.22-μm pore size membrane. Protein concentration of the supernatant (SEA) was determined by the Bradford Protein Assay (80) and aliquots of the SEA stored at -80°C.

### Guide RNAs, Cas9, and single stranded DNA repair template

Single guide RNA (sgRNA) was designed using the web-based tools at http://bioinfogp.cnb.csic.es/tools/breakingcas/(81) to predict cleavage sites for the *Streptococcus pyogenes* Cas9 nuclease within the genome of *S. mansoni*. The sgRNA targeted exon 6 of the ω1 gene, Smp_193860, www.genedb.org, residues 3808-3827, adjacent to the protospacer adjacent motif, AGG (Fig. 1a). This is a multi-copy gene with at least five copies of ω1 located in tandem on chromosome 1 (12). To infer the gene structure of Smp_193860 in the *S. mansoni* V5 assembly more accurately, the omega-1 mRNA DQ013207.1 sequenced by Fitzsimmons *et al*. (2005) (29) was used to predict the gene structure with the exonerate software, by aligning it to the assembly using the exonerate options ‘--model coding2genome’ and ‘--maxintron 1500’. The Smp_193860 copy of ω1 includes nine exons interspersed with eight introns (6,196 nt) (Fig. 1a).

Synthetic gRNA (sgRNA), ω1-sgRNA was purchased from Thermo Fisher Scientific (Waltham, MA). A double stranded DNA sequence complementary to the sgRNA was inserted into lentiviral gene editing vector, pLV-U6g-EPCG (Sigma), which encodes Cas9 from *S. pyogenes* driven by the eukaryotic (human) translation elongation factor 1 alpha 1 (tEF1) promoter and the sgRNA driven by the human U6 promoter (Fig 1c). (The pLV-U6g-EPCG vector is tri-cistronic and encodes the reporter genes encoding puroR and GFP, in addition to Cas9 (82).) This gene-editing construct, targeting exon 6 of ω1 Smp_193860, was termed pLV-ω1X1. A single stranded oligodeoxynucleotide (ssODN), which included homology arms of 50 nt each in length at the 3’ (position 3774-3824 nt) and 5’ (3825-3874 nt) flanks and a small transgene (5’-<u>TAA</u>G<u>TGA</u>C<u>TAG</u>G<u>TAA</u>C<u>TGA</u>G<u>TAG</u>-3’, encoding stop codons (six) in all open-reading frames) (Fig. 1b), was synthesized by Eurofin Genomics (KY, USA). An oligonucelotide primer that included this sequence was employed in PCRs to investigate the presence of CRISPR/Cas9-programmed insertion of the transgene (83) (Table s2).

### Transfection of schistosome eggs with a Cas9/guide RNA complex

For the ribonucleoprotein (RNP) complex of the ω1-sgRNA and recombinant Cas 9 from *Streptococcus pyogenes*, 3 μg of ω1-sgRNA and 3 μg of Cas9-NLS nuclease (Dharmacon, Lafayette, CO) were mixed in 100 μl Opti-MEM (Sigma) to provide 1:1 ratio w/w RNP. The mixture was incubated at room temperature for 10 min, pipetted into a 4 mm pre-chilled electroporation cuvette containing ∼10,000 LE in ∼150 μl Opti-MEM, subjected to square wave electroporation (one pulse of 125 volts, 20 milliseconds) (BTX ElectroSquarePorator, ECM830, San Diego, CA). The electroporated eggs were incubated for 5 min at room temperature, and maintained at 37°C, 5% CO_2_ in air for 3, 5 and 7 days. To investigate whether homology-directed repair (HDR) could catalyze the insertion of a donor repair template, 3 μg ssODN was mixed with RNP and the LE before electroporation. In a second approach (above), the ssODN was delivered to LE by electroporation at ∼24 hours after the lentiviral transduction of the LE. The eggs were collected 3, 5 and 7 days later and genomic DNA recovered from LE. The negative controls included LE subjected to electroporation in the presence of only Opti-MEM, only Cas 9, only sgRNA, and only ssODN.

### Transduction of schistosome eggs with lentiviral particles

*Escherichia coli* Zymo 5α (Zymo Research) cells were transformed with lentiviral plasmid pLV-ω1X1 and cultured in LB broth in 100 μg/ml ampicillin at 37°C, agitated at 225 rpm for ∼18 hours, after which plasmid DNA was recovered (GenElute Plasmid purification kit, Invitrogen). A lentiviral (LV) packaging kit (MISSION, Sigma-Aldrich) was used to prepare LV particles in producer cells (human 293T cell line). In brief, 3.5 × 10^5^ of 293T cells/well were seeded in a 6-well tissue culture plate in DMEM supplemented with 10% heat-inactivated fetal bovine serum (FBS), 2 mM L-glutamine, 1% penicillin/streptomycin and cultured at 37°C, 5% CO_2_ for 18 hours. The producer cells were transfected using FUGENE HD (Promega) with pLV-ω1X1 and LV packaging mix containing two additional plasmids; one plasmid that expressed HIV structural and packaging genes and another that expressed the pseudotyping envelope protein Vesicular Stomatitis Virus Glycoprotein (VSVG). Subsequently, the transfection mixture (50 μl; 500 ng plasmid DNA, 4.6 μl packaging mix, 2.7 μl of FUGENE HD in Opti-MEM) was dispensed drop wise into each well on the plate. Sixteen hours later, the media were removed from the transfected cells, replaced with pre-warmed complete DMEM, and cells cultured for 24 hours. The supernatant, containing VSVG-pseudotyped LV particles was filtered through 22 μm pore size membranes (45), and stored at 4°C. Additional pre-warmed complete DMEM was added to the well, for culture for a further 24 hours. The supernatant was collected as above, combined with the first supernatant and concentrated (Lenti-X concentrator, Takara Bio, Mountain View, CA). Virion titer was estimated by two methods; first, by use of Lenti-X-GoStix (Takara Bio) to establish the presence of functional virions at >10^5^ infectious units (IFU)/ml, and second, by reverse transcriptase assay (45, 84) to quantify levels of active virions. Virions with counts of ∼4×10^6^ count per minute (cpm)/ml were aliquoted and stored at -80°C.

To transduce LE with LV, ∼10,000 eggs were incubated for 24 hours in complete DMEM containing 500 μl of ∼4×10^6^ cpm/ml VSVG-LV virions. Thereafter, the LE were washed three times in 1×PBS and transfected with ssODN by square wave electroporation or further steps. LV virions heat-inactivated by incubation at 70°C for two hours (45) with subsequent transfection with the ssODN, transfection with ssODN in the absence of virions or Opti-MEM only served as negative controls.

### PCR amplification of diagnostic transgene to detect knock-in into exon 6 of ω1

For each DNA sample, four separate PCR assays using four distinct primer pairs (Table s2) were carried out. The first ω1 primer pair, to amplify locations 3751-4740 nt of Smp_193860, was employed as positive control for the presence of genomic DNA with the Smp_193860 copy of *ω1*. The other primer pairs shared one forward primer complementary to the knock-in 24 nt transgene with three reverse primers, Sm ω1-R1, -R2 and -R3 at positions 3966-3984, 4066-4085 and 4102-4121 nt, respectively, binding to three sites downstream of the ω1 predicted DSB site (Fig. 2a; Table s2) (83). The PCR mix included 10 μl Green GoTaq DNA polymerase mix (Promega) with 200 nM of each primer and 10 ng genomic DNA. Thermal cycling conditions involved denaturation at 95°C, 3 min followed by 30 cycles of 94°C, 30 seconds, 60°C, 30 seconds and 72°C, 30 seconds and a final extension at 72°C for 5 minutes. Following agarose gel electrophoresis (1.2% agarose/TAE), amplicons of the expected sizes were recovered from gels and ligated into pCR4-TOPO (Thermo Fisher). *E. coli* Zymo 5α competent cells were transformed with the ligation products, several colonies of each transformant were grown under ampicillin selection, plasmid DNA purified, and insert sequenced to confirm the presence and knock-in of the transgene (Fig. 1c).

### Illumina sequencing

Pooled LE DNA samples from six independent KI experiments of pLV-ω1X1 with ssODN were used as the template to amplify the on-target DNA fragment using MiSeq primers (Fig. 2a) with High Fidelity *Taq* DNA polymerase (Thermo Fisher). PCR reactions were set up with 10 ng LE DNA samples from the KI experiment in 25 µl reaction mix using the HiFidelity Taq DNA polymerase (Thermo Fisher) following the PCR program 94ºC for 3 minutes of denaturation followed by 30 cycles of 94ºC for 30 seconds, 60ºC or 54ºC for 30 seconds, 72ºC for 45 seconds and final extension at 72ºC for 2 minutes. The expected size of the amplicon flanking predicted DSB was 205 bp. Amplicons of this size were purified using the Agencourt AMPure XP system (Beckman Coulter). Amplicons generated from four different PCR reactions from each sample were pooled, and 100 ng of amplicons from each sample was used to construct the uniquely indexed paired-end read libraries using the QIAseq 1-step Amplicon Library Kit (Qiagen) and GeneRead Adapter I set A 12-plex (Qiagen). These libraries were pooled, and the library pool was quantified using the QIAseq Library Quant System (Qiagen).

Samples (Table s3) were multiplexed (10 samples) and each run on four MiSeq lanes. After sequencing, the fastq files for each particular sample were merged. Samples 1-6, 8 and 10 were prepared using an annealing temperature of 54ºC. Samples 7 and 9 were prepared using an annealing temperature of 60ºC, and included an extra 10 bp at the start of the MiSeq sequences, ‘GTTTTAGGTC’, present upstream of the 5’ primer in the genomic DNA. We trimmed this sequence from the reads using cutadapt v1.13 (85). To detect HDR events, CRISPResso v1.0.9 (48, 86) was employed using a window size of 500 bp (-w 500) with the reference amplicon according to gene *Smp_193860* in the *S. mansoni* V7 assembly, and with the --exclude_bp_from_left 25 and --exclude_bp_from_right 25 options in order to disregard the (24 bp) primer regions on each end of the amplicon when indels are being quantified. A window size of 500 nt was employed to include the entire amplicon. In order to search for HDR events, CRISPResso checked for HDR events (using –e and –d options) in treatment groups including controls. To infer frameshifts using CRISPResso the –c option was used, giving CRISPResso the coding sequence from positions 42-179 of the amplicon. To confirm the insertions of the knock-in sequences reported by CRISPResso (column L in Table s3), we took all insertions of 20-28 bp reported by CRISPResso, and calculated their percent identity to the expected knock-in sequence using ggsearch v36.3.5e in the fasta package (87), and an insertion was considered confirmed if it shared ≥75% identity to the expected donor knock-in sequence.

### Copy number estimation for *ω1*

A quantitative PCR to estimate the relative copy number of ω1 was performed using Kapa qPCR mastermix SYBRfast (KK4602) on 1 ng of gDNA templates isolated from control and test samples, in 20 μl volumes. Primer pair OMGgRNA1F and OMGgRNA1R was used to amplify the ω1 gRNA target region and SmGAPDH (Smp_056970) as a reference single-copy gene (primers shown in Table s2). The PCR efficiencies for primer pairs were estimated by titration analysis to be 100% ±5 (88) and qPCRs were performed in triplicate in 96-well plates, with a denaturation step at 95°C of 3 min followed by 40 cycles of 30 sec at 95°C and 30 sec at 55°C, in thermal cycler fitted with a real time detector (StepOnePlus, Applied Biosystem). The relative quantification assay 2^−ΔΔCt^ method (89) was used to ascertain the relative copy number of *ω1.* Relative copy number of *ω1* in the CRISPR/Cas9 treated groups reflects the fold change of *ω1* copy number normalized to the reference gene (Sm GAPDH) and relative to the untreated control group (calibrator sample with relative *ω1* copy number = 1) (49).

### Gene expression for *ω1* mRNA

Total RNAs from schistosome eggs were extracted using the RNAzol RT reagent (Molecular Research Center, Inc), which eliminates contaminating DNA (90), and concentration and purity determined using a spectrophotometer (OD_260/280_ ∼2.0). Reverse transcription (RT) of the RNA (500 ng) was performed using iScript Reverse Transcript (Bio-Rad), after which first strand cDNA was employed as template for qPCRs using SsoAdvanced Universal SYBR Green Supermix (Bio-Rad) performed in triplicates in an iQ5 real time thermal cycler (Bio-Rad). RT-qPCR reaction mixtures included 2 μl first strand cDNA, 5 μl SsoAdvanced Universal SYBR Green Supermix, and 300 nM schistosome gene specific primers. Table s2 provides details of the oligonucleotide primers. Thermal cycling included denaturation at 95°C for 30 sec, 40 amplification cycles each consisting of denaturation at 95°C for 15 sec and annealing/extension at 60°C for 30 sec, and a final melting curve. The output was analyzed using the iQ5 software. Relative expression was calculated using the 2^−ΔΔCt^ method and normalized to schistosome GAPDH expression (89); data are presented as transcript levels (three replicates) compared to the WT (100%) LE, and fold change reported as mean relative expression ± SD.

### Nuclease activity of ω1

A stock solution of yeast RNA (Omega Bio-tek, Norcross, GA) was prepared at 1.0 μg/μl, 50 mM Tris-HCl, 50 mM NaCl, pH 7.0. Yeast RNA (200 ng) was incubated with 2 μg SEA from control and experimental groups individually at 37°C for 60 min. (SEA investigated here, named Δω1-SEA, was extracted from LE transduced with pLV-ω1X1 virions and ssODN, pooled from six biological replicates.) RNase A, an endoribonuclease from bovine pancreas (Thermo Fisher) served as a positive control enzyme whereas yeast RNA in reaction buffer only served as the negative control. The RNase activity of ω1 in wild type SEA or Δω1-SEA was analyzed by visualizing and quantifying the substrate that remained following enzymolysis by agarose gel electrophoresis and staining with ethidium bromide. The yeast RNA digestion by control SEAs or Δω1-SEA were set up in triplicates, with quantity of residual RNA determined by densitometry (50).

### Macrophage polarization by WT or Δω1-SEA and T-cell activation *in vitro*

Human monocytic THP-1 cells were maintained in Roswell Park Memorial Institute medium (RPMI) 1640 with L-glutamine, HEPES (Thermo Fisher Scientific) containing 10% (v/v) FBS with 4 mM glutamine, 25 mM HEPES, 2.5 g/L D-glucose at 37°C in 5% CO_2_ in air. THP-1 cells were non-adherent cells. In a 6-well plate, THP-1 monocytes (3×10^5^ cells in each well) were differentiated into macrophages (Mϕ) by incubation in 150 nM phorbol 12-myristate 13-acetate (PMA) (Sigma) for 48 hours (91). Mϕ were exposed to SEA (50 ng/ml) or Δω1-SEA (50 ng/ml) (from LE transduced with pLV-ω1X1 virions and ssODN) for 48 hours. To investigate macrophage and T cell interactions, Mϕ cells were pulsed with 50 ng/ml SEA or Δω1-SEA and thereafter co-cultured in direct contact with Jurkat (human CD4^+^ T) cells. Nine ×10^5^ Jurkat were added to Mϕ with direct contactand were co-cultured for an additional 72 hours. Cell-free supernatants from the co-cultures were collected to quantify secretion of T helper cell cytokines including IL-4, IL-5, IL-13, IL-10, TNF-α, IL-6, IL-2 and IFN-γ by enzyme linked immunosorbent assay (ELISA) (Qiagen) (92). The assay included positive controls for each analyte, which were provided in the ELISA kit (Qiagen). Three biological replicates were undertaken.

### Schistosome egg-induced primary pulmonary granulomas

For induction of circumoval, egg-induced granulomas in the lungs of mice, 8 week old female (25) BALB/c mice were injected with 3,000 WT eggs or Δω1-eggs (from experiment pLV-ω1X1 with ssODN) or 1×PBS as negative control by tail vein, as described (58). The mice (6-9 mice/group) were euthanized 10 days later. For histopathological assessment of granuloma formation, the left lung was removed at necropsy and fixed in 10% formalin in pH 7.4 buffered saline for 24 hours, after which it was dehydrated in 70% ethanol, and clarified in xylene. The fixed lung tissue was embedded in paraffin and sectioned at 4-µm-thickness by microtome (58). Thin sections of the left lung lobe were mounted on glass slides and fixed at 58-60°C. Subsequently, rehydrated sections were stained with hematoxylin-eosin (H&E) for evaluation of inflammatory infiltrates and cellularity of granulomas. The longest (R) and shortest (r) diameters of each granuloma containing a single egg were measured with an ocular micrometer, and the volume of the granuloma calculated assuming a prolate spheroidal shape, using 4/3 π Rr^2^ (25). All granulomas in all slides from the left lung of the mice, 15 slides per treatment group, were measured; in total, >100 granulomas from each treatment group. Digital images were captured using a 2D glass slide digital scanner (Aperio Slide Scanner, Leica Biosystems, Vista, CA) and examined at high magnification using the Aperio ImageScope (Leica) software (57, 93).

### Statistics

Means for experimental groups were compared to controls by one-way ANOVA and, where appropriate, by two-tailed Student’s *t*-test and Welch’s unequal variances *t*-test (GraphPad Prism, La Jolla, CA). Values for *p* of ≤ 0.05 were considered to be statistically significant.

## Acknowledgements

We thank Dragana Jankovic, Alan Sher, Thomas Wynn, Michael Bukrinsky, Larisa Dubrovsky, Arnon Jurberg, Meredith Brindley, Robert Thompson and Thiago De Almeida Pereira for advice and technical assistance. Schistosome-infected mice and snails were provided by the NIAID Schistosomiasis Resource Center of the Biomedical Research Institute, Rockville, Maryland through NIH-NIAID Contract HHSN272201000005I for distribution through BEI Resources. We acknowledge the Thailand Research Fund through Senior Research Scholar Grant (RR, W. Maleewong) and the Royal Golden Jubilee Ph.D. program numbers PHD/0011/2555 (AC, S. Pinlaor), PHD/0047/2556 (PP, Nitat Sookrung), PHD/0053/2556 (RR, W. Maleewong). These studies were supported by Wellcome Trust Strategic Award number 107475/Z/15/Z (KFH, principal investigator), Wellcome Trust grant WT 098051 award (MB), R21AI109532 (GR, PJB) award from NIAID, National Institutes of Health, and, in part, by generous support from MaxMind Inc./TJ Mather (PJB).

## Supporting Information

## Supplementary Tables

**Table s1**. Putative NHEJ and HDR pathway genes in *S. mansoni*.

**Table s2**. List of oligonucleotide primers. Nucleotide sequences and position on the schistosome *omega-1* gene; Smp_193860.

**Table s3**. Frequency of knock-in sequences, indels and substitutions in Illumina sequencing data, considering the entire 202-bp amplicon (except for 25-bp at the ends, to exclude primer regions)

## Supplementary methods

### Estimation of mutation efficiency

We used the egg genomic DNA templates directly for qPCR as described (46, 47), with slight modification which enabled estimation of the efficiency of CRISPR-mediated mutagenesis at the target locus without the need to normalize the experimental and control template DNAs.

The general approach makes use of the fact that the binding of a primer overlapping the sgRNA site was compromised in programmed mutagenized egg (LE) genome(s), resulting in delayed amplification, whereas binding of a flanking primer pair was unaffected. The ‘OUT’ (flanking) primer pair encompassed at least 50 bp surrounding the sgRNA binding region. The ‘OVR’ (overlapping) primer pair used one of the OUT primers and another primer that bound the 20 bp of the sgRNA target sequence. The 3′ side of the OVR primer bound immediately upstream of the NGG motif, as the majority of indels would be expected to affect positions −1 to −10 of the binding site (86)(Fig. 1; Fig. s5).

At days 3, 5 and 7 following transfections with or without ssODN, genomic DNA was isolated from LE. Using 5 ng of DNA template, separate 20 µl OUT and OVR qPCR reactions were undertaken. Quantitative PCR was performed with SsoAdvanced SYBR Green Supermix (Bio-Rad, 172-5271) using a Bio-Rad iQ5 Real-Time PCR system, with qPCR conditions at initial 95°C for 30 s, 40 cycles, 95°C for 10 s, 60° C for 20 s. Primers are listed in Table s2.

The ratio of the qPCR quantification cycle values for the control OVR and control OUT primers reflected the differences in amplification of the two primer pairs on control DNA template. This might have been due to inherent differences in amplification that exist even between perfectly complementary primer pairs. In contrast, the OVR/OUT ratio in mutant DNA reflected both this difference in amplification between the primer pairs and the loss of the OVR binding site due to CRISPR-introduced indels. A comparison of the OVR/OUT quantification cycle ratios of control versus mutated genomes thus reflected the efficiency of mutagenesis.

The CRISPR efficiency was calculated by the Ct ratio of OVR:OUT, after which indel/substitution mutation percentage was estimated as follows:

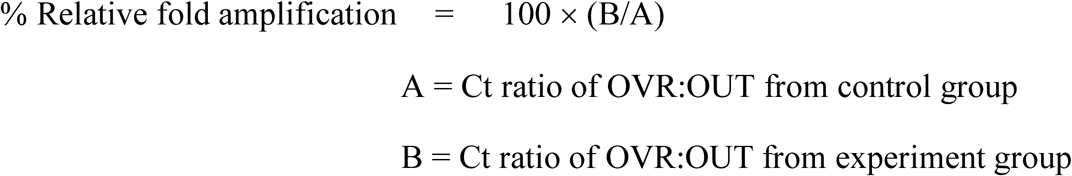

ω1X1 virions (green), pLV-ω1X1 virions and ssODN (blue) compared to controls. **c,** Loss of RNase activity as assessed by hydrolysis of yeast RNA. Residual yeast following exposure to SEA, visualized after gel electrophoresis; lane 1, buffer (negative control); 2, RNase A; 3-8, WT SEA and other control SEAs as indicated in D; 9-12, Δω1-SEA from RNP, RNP and ssODN, pLV-ω1X1 virions, and pLV-ω1X1 virions and ssODN, respectively. **d,** Intact yeast RNA (nanograms) remaining following incubation with SEA (mean ±SD, *n* = 6). More RNA remained following incubation with Δω1-SEA in all groups, RNP, RNP and ssODN, pLV-ω1X1 virions, and pLV-ω1X1 virions and ssODN treated SEA) (blue) than in the WT SEA controls (*p* ≤ 0.0001). Among the gene edited experimental groups, more RNA remained when donor template was introduced at the same time as RNP or pLV-ω1X1 virions (*p* ≤ 0.01). Significant differences were not apparent among the WT SEA control groups.

**Figure s1.**
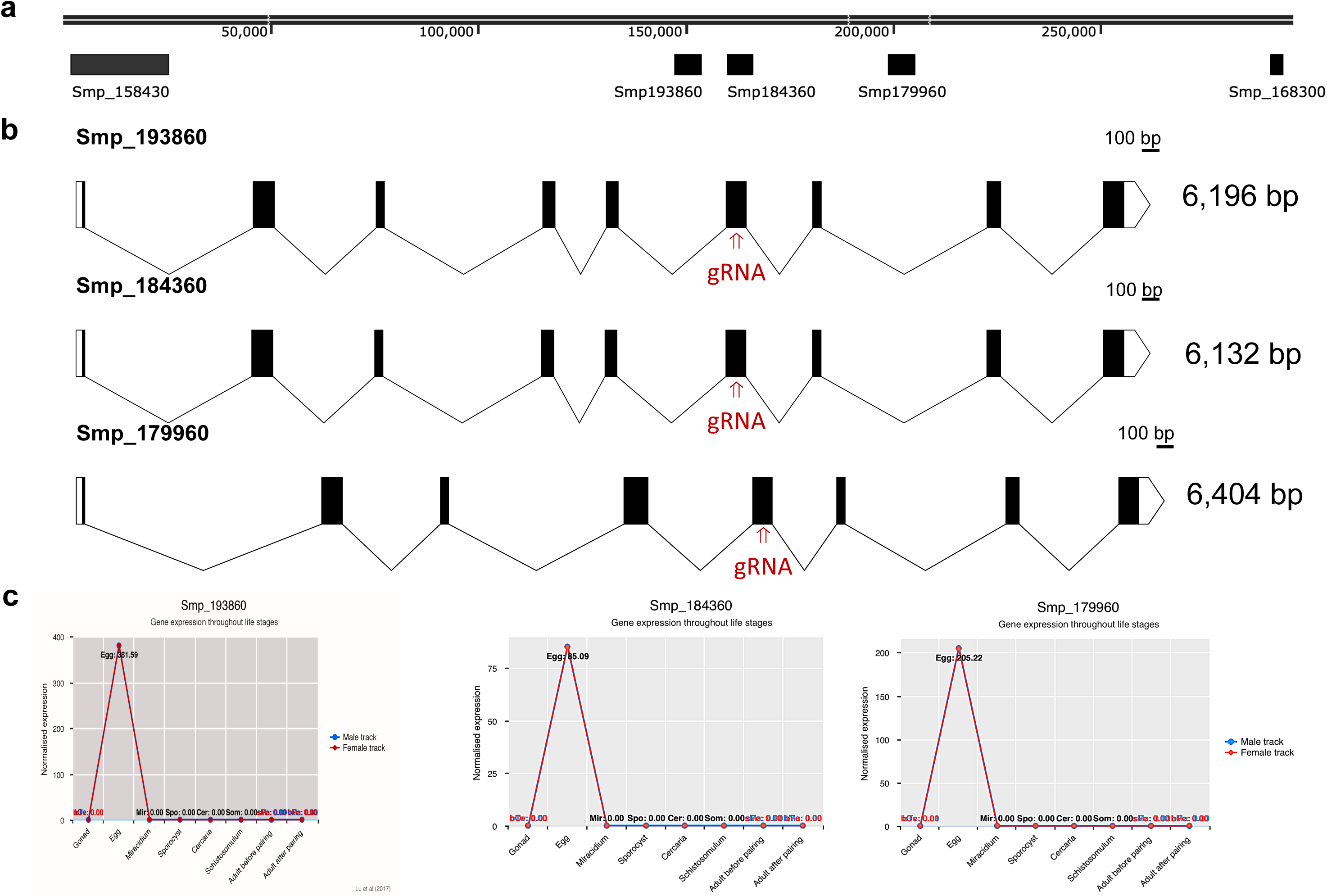
The T2 ribonuclease ω1 is encoded by at least five gene copies located on schistosome chromosome 1. Panel **a**. The position of several genes coding for ω1, including *Smp_179960, Smp_184360, Smp_193860* are shown, to indicate the physical relationship of the copies on the chromosome. Two additional copies of ω1, Smp_158430 and Smp_168300 do not include the cognate sgRNA and PAM motifs. **b,** Map of gene structure and size of *Smp_179960, Smp_184360, Smp_193860*, along with the ω*1*-specific sgRNA target site. **c,** Normalized expression of three copies of ω1 as indicated throughout the developmental stages of *S. mansoni*. The profile of developmental expression of ω1 was established by meta-analysis of RNAseq reads (73), https://doi.org/10.1101/308213

**Figure s2.**
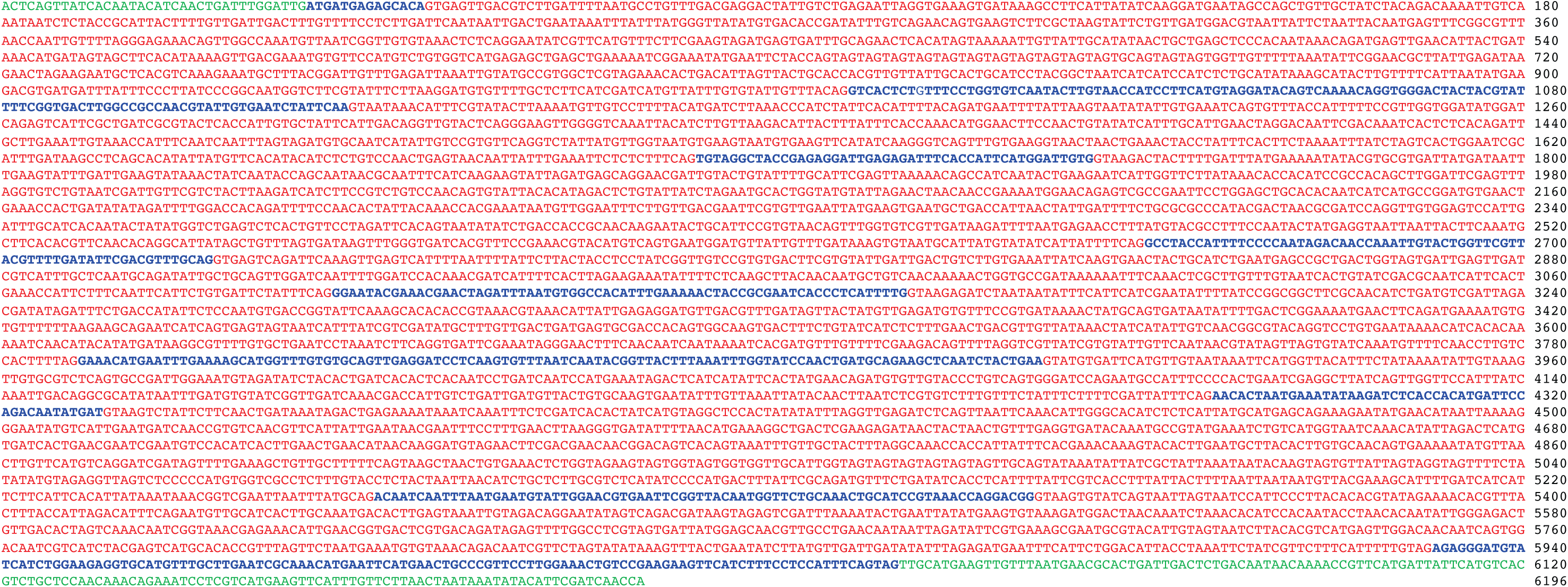
The nucleotide sequence of the Smp_193860 copy, and indicates the UTR (green), coding exons (blue) and introns (red).

**Figure S3.**
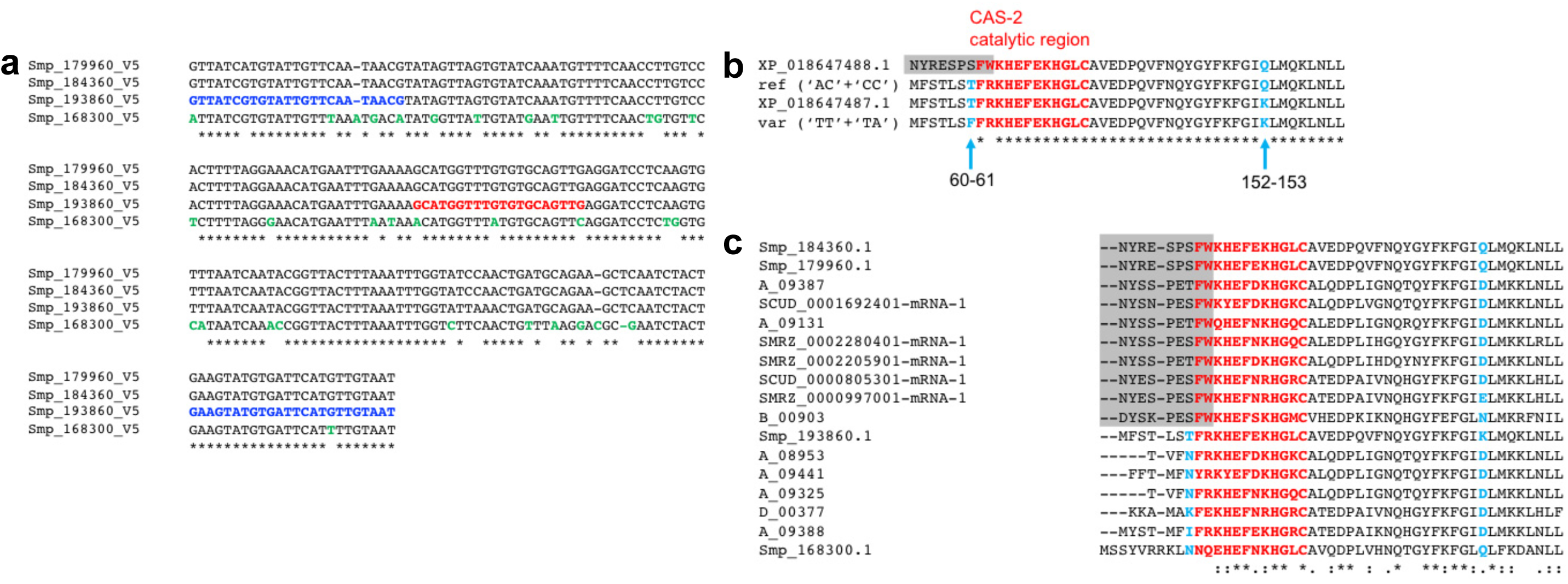
Variation among copies of *ω1* in the reference genome. Panel **a**, An alignment assembled using CLUSTALW (97) of the MiSeq amplicon region in the ω*1* genes *Smp_179960, Smp_184360, Smp_193860* and *Smp_168300* using the sequences from the *S. mansoni* genome, assembly V5. The MiSeq primers (highlighted in blue) and sgRNA (red) were designed based on *Smp_184360*. Residues highlighted in green show where *Smp_168300* differs from the other three genes. **b**, An alignment made using CLUSTALW (97) of the coding region of the amplicon sequence found in the gene *Smp_193860* in the *S. mansoni* V7 assembly, the predicted alternative sequence for the variant having ‘TT’ at position 60-61 and ‘TA’ at 152-153 in the amplicon (‘var’), and the translations of two mDNAs retrieved from NCBI (top BLASTP hits in the NCBI protein database). Note that this corresponds to a part only of the ω**1** T2 ribonuclease. The CAS-2 catalytic region is highlighted in red (29). The mRNA XP_018647488.1 has an alternative splicing event at the start of the exon included in the amplicon (shaded grey), so that it only matches the region from ‘KHEFEK…’ (in the CAS-2 catalytic region) onwards. **c**, An alignment of the coding region of the amplicon sequence in schistosome ω1 homologues (family 873078) (40), excluding diverged members from *S. japonicum* as well as the diverged *S. haematobium* gene *B_00112*. The region with sequence similarity to the alternative splice-form is in shaded grey. Note that the *Smp_193860* sequence used in this alignment is the version in the *S. mansoni* V5 gene set, which has the Q->K substitution at 152-153.

**Figure s4.**
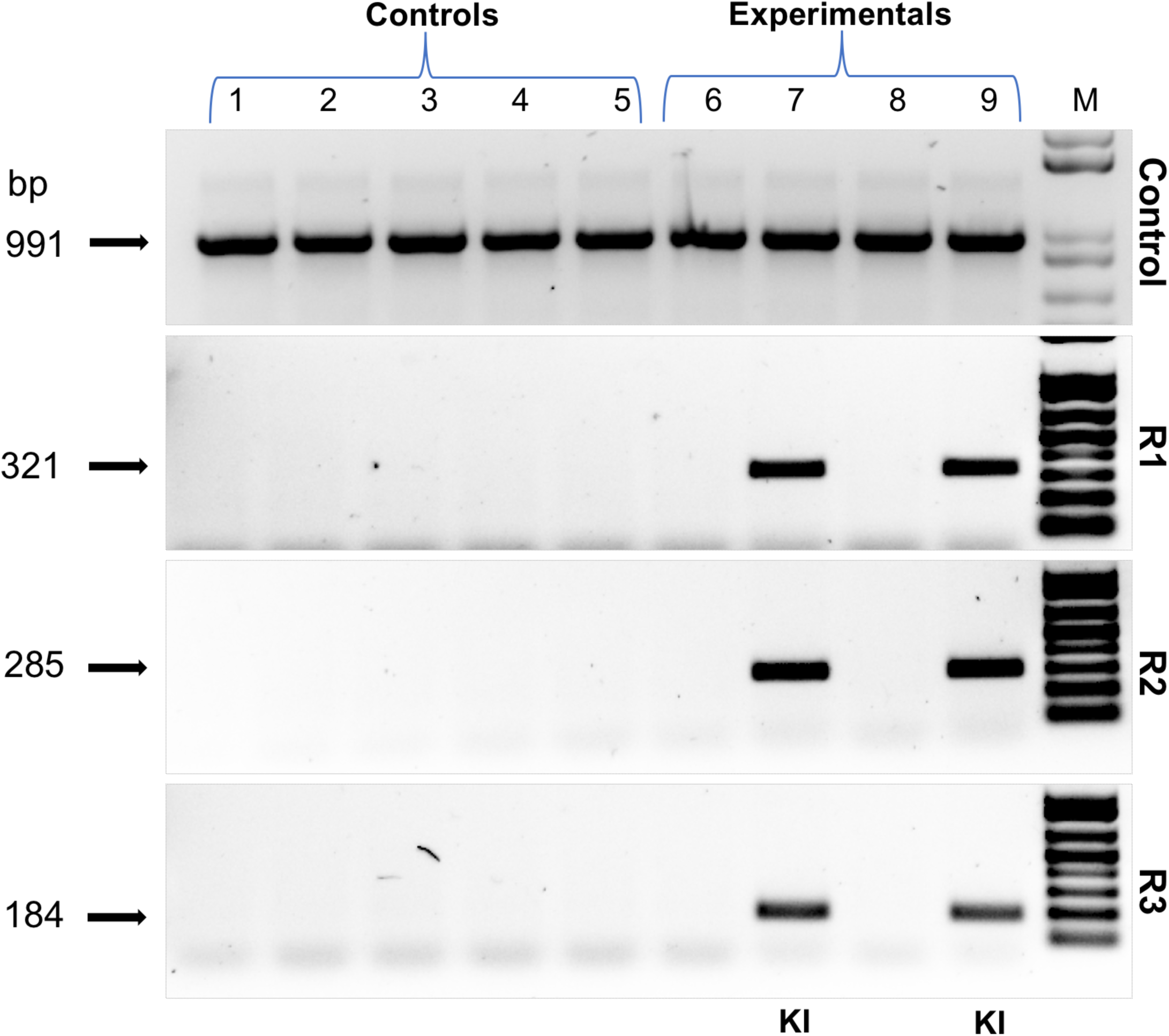
Programmed HDR-catalyzed knock-in of transgene into ω1, exon 6. The PCR products visualized in ethidium bromide-stained agarose gel demonstrated Cas9 catalyzed target site-specific insertional mutagenesis in exon 6 of the ω1 gene. The 24 nt transgene knocked-in into target site indicated in lanes R1, R2 and R3 of 321, 285 and 184 bp, respectively (arrows at left) spanning the mutated site in the genomic DNAs pooled from schistosome eggs, including a positive control flanking the insert site (991 bp). Lanes 1-5 show the amplicons from the control groups; medium only, sgRNA only, cas9 only, ssODN only and heat-inactivated pLV-ω1X1+ssODN. Lanes 6-9 show the amplicons from experiment groups; RNP, RNP and ssODN (KI), pLV-ω1X1virions, and pLV-ω1X1virions and ssODN (KI). Programmed knock-in was detected only in the KI experimental groups, as shown in lanes 7 and 9 where the transgene-specific amplicon obtained with the R1, R2 and R3 primers was present. This product was not seen in PCRs where DNAs from control and other experimental groups were amplified. Amplicons of 991 bp were seen with all genomic DNAs, confirming the integrity of the genomic DNAs and the control amplicon primer pair.

**Figure s5.**
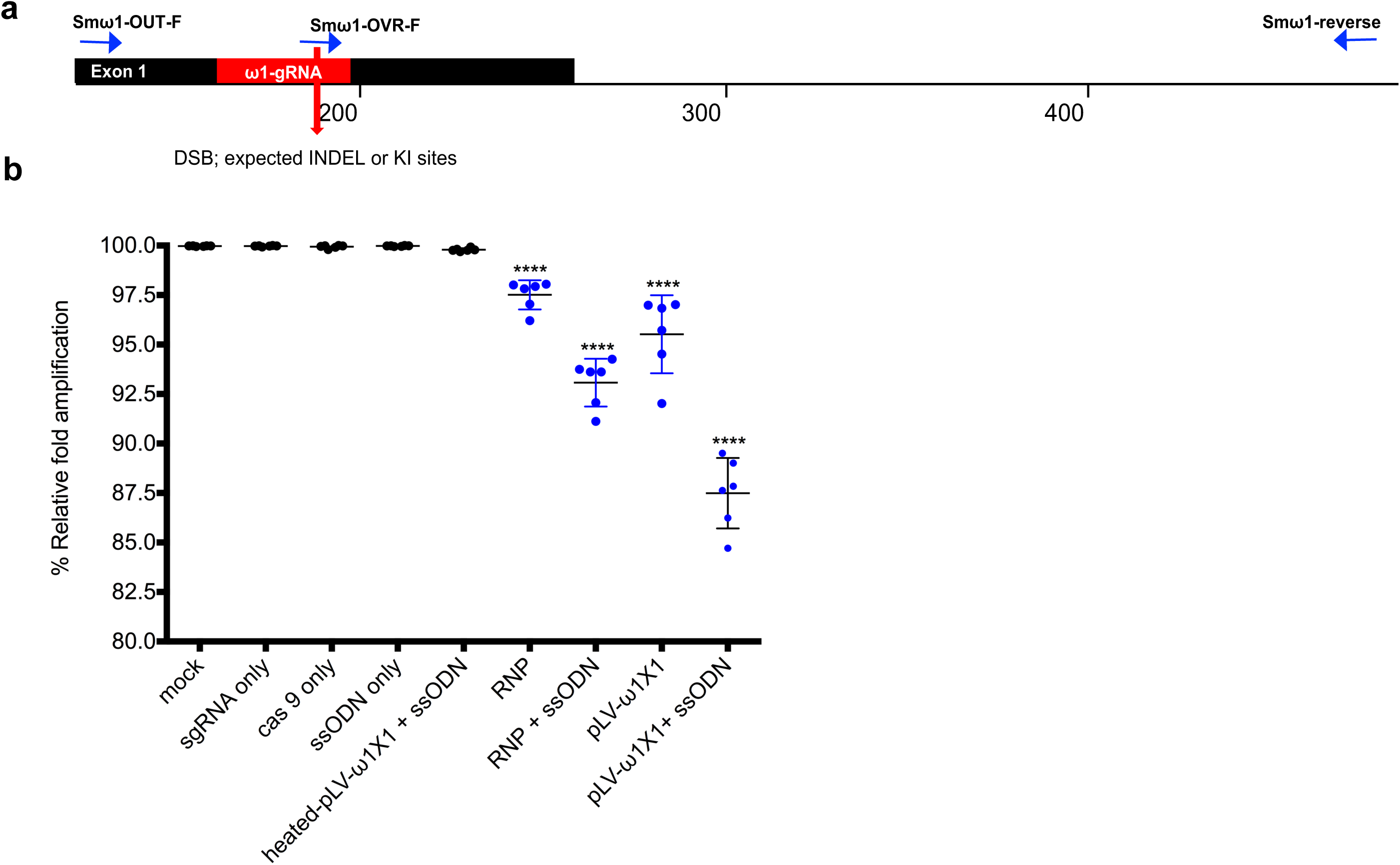
Quantitative PCR to estimate efficiency of programmed mutagenesis. Panel **a.** The three primers (tri-primer set): Smω1-OVR-F (probe at gRNA, expected DSB and KI sites as red bar and arrow), Smω1-OUT-F and shared Smω1-reverse primer locations used in SYBR green-based PCR to estimate efficiency of on-target, programmed CRISPR/Cas9 gene editing. Estimation of gene-editing efficiency was determined as described (46, 47). **b,** Relative fold amplification efficiency; the ratio of OVR:OUT (gray bar) was set at 100% for control DNA sample, and little or no reduction was observed in relative fold amplification compared to other control groups, including groups exposed to culture medium only, sgRNA only, Cas9 only, ssODN only, heat-killed pLV-ω1X1 virions and ssODN. By contrast, there was reduced relative fold amplification of 2.5%, 6.9%, 4.5% and 12.5% in the experimental groups, RNP, RNP and ssODN, pLV-ω1X1 virions, and pLV-ω1X1virions and ssODN, respectively. Percentages of relative fold amplification differed significantly among control and experiment groups. Means ± SE were established from six biological replicates.

**Figure s6.**
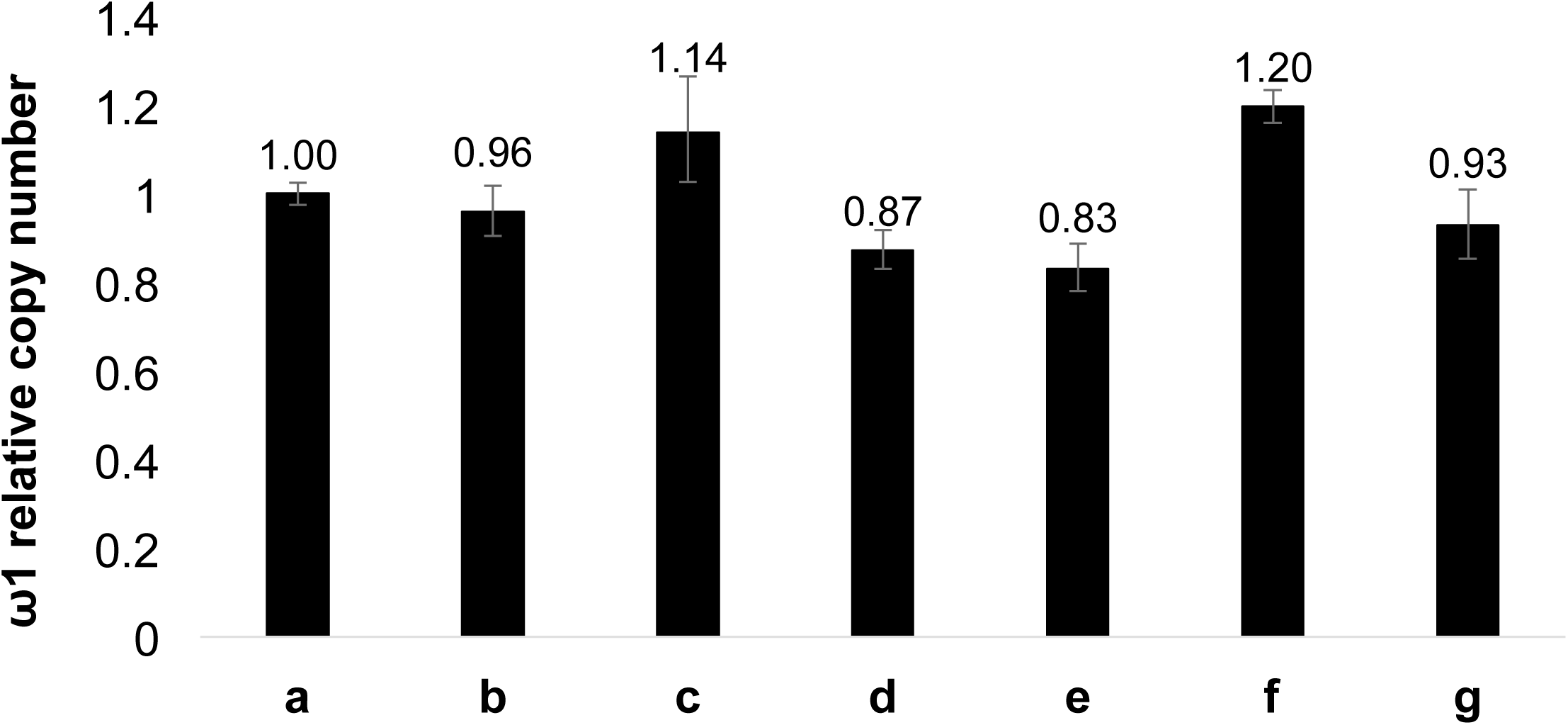
Estimated relative copy number of *ω1*. Copy number estimated by qPCR in genomic DNAs isolated from CRISPR/Cas9-treated eggs or controls. The relative copy number of the calibrator sample a = 1. Means ±SD: a, eggs; b, eggs and Cas9; c, eggs and ssODN; d, eggs and heat-inactivated pLV-ω1X1 virions; e, eggs, heat-inactivated pLV-ω1X1 virions and ssODN; f, eggs and pLV-ω1X1 virions; g, eggs, pLV-ω1X1 virions and ssODN.

**Figure s7.**
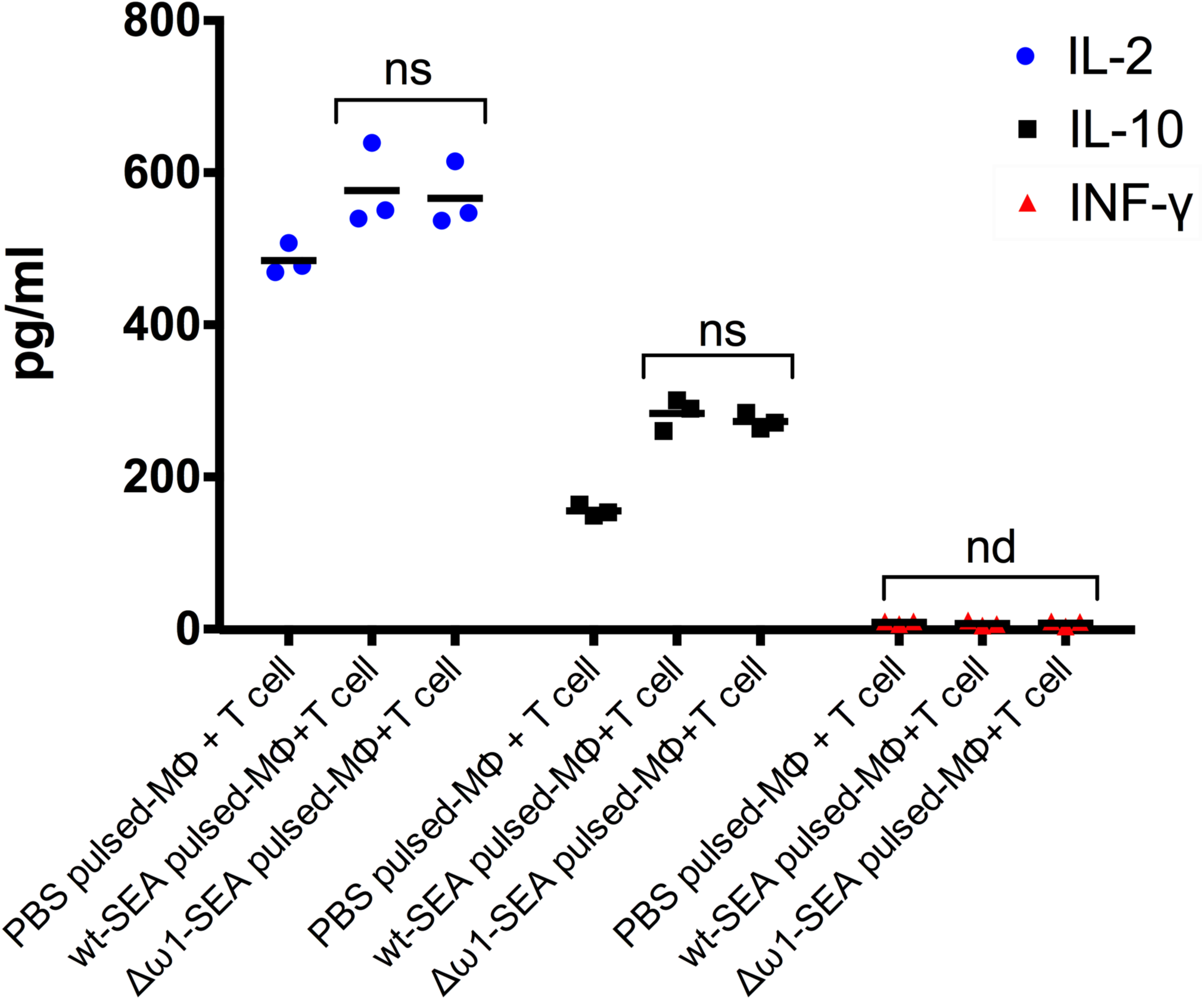
Levels of IL-2, IL-10 and IFN-γfollowing exposure to Δω1-SEA. Levels of IL-2 (blue circle) and IL-10 (black square) were not significantly different among groups after pulsing the Mϕ (PMA-induced THP-1 cells) with Δω1-SEA prior to co-culture with CD4^+^ T cells compared with WT-SEA pulsed-Mϕ. IFN-γ (red triangle) was not detected in any of the three groups.

**Table s1a.**
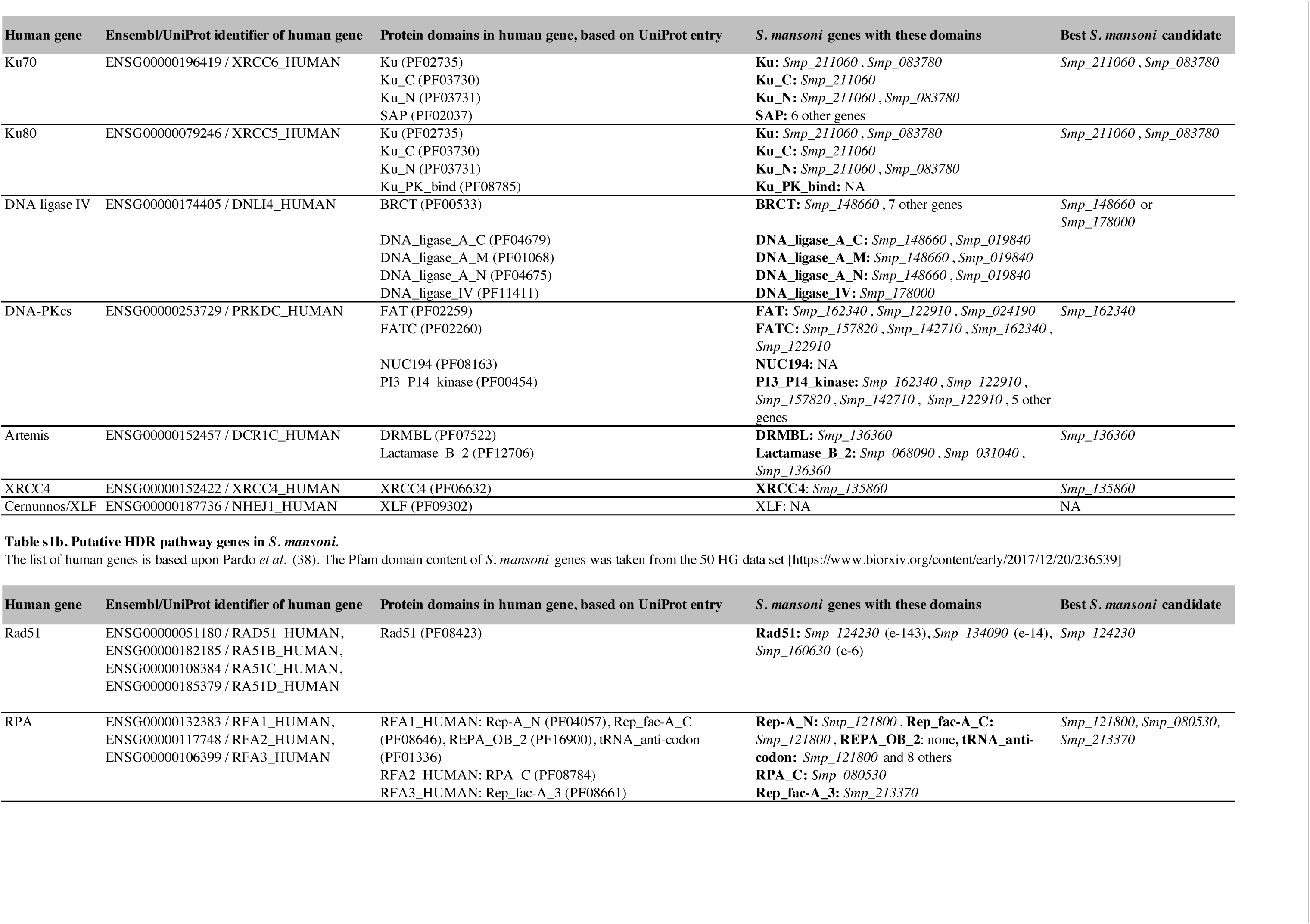
Putative NHEJ pathway genes in *S. mansoni.* The list of human genes taken from Table 1 in Lee *et al.* (36). The Pfam domain content of *S. mansoni* genes was taken from the 50 HG data set [https://www.biorxiv.org/content/early/2017/12/20/236539]

**Table s2.**
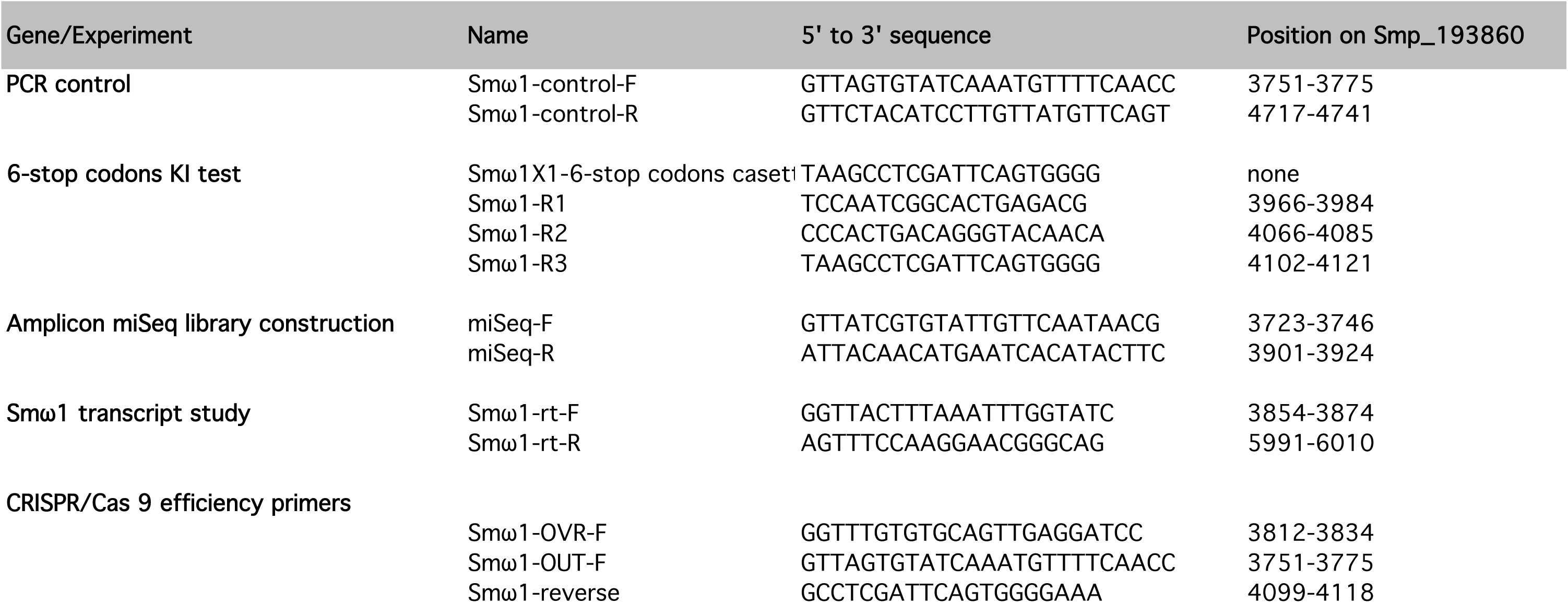
Primer sequences

**Table s3.**
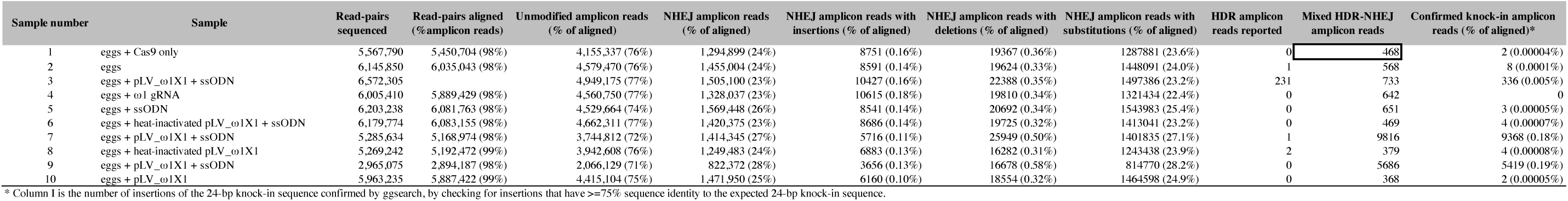
Frequency of knock-in sequences, indels and substitutions in MiSeq sequencing data. The CRISPResso analysis used a window size (-w option) that included the whole 202 bp amplicon, except 25 bp at each end in order to exclude the primer regions.

